# Organ-specificity of sterol and triterpene accumulation in *Arabidopsis thaliana*

**DOI:** 10.1101/2020.03.23.004358

**Authors:** B. Markus Lange, Brenton C. Poirier, Iris Lange, Richard Schumaker, Rigoberto Rios-Estepa

## Abstract

Sterols serve essential functions as membrane constituents and hormones (brassinosteroids) in plants, while non-sterol triterpenoids have been implicated in defense responses. Surprisingly little is known about the sterol and triterpene profiles in different plant organs. To enhance our understanding of organ-specific sterol and triterpene accumulation, we quantified these metabolite classes in four different organs (root, stem, leaf, seed) of *Arabidopsis thaliana* (L.). Based on these data sets we developed kinetic mathematical models of sterol biosynthesis to capture flux distribution and pathway regulation in different organs. Simulations indicated that an increased flux through the sterol pathway would not only result in an increase of sterol end products but also a concomitant build-up of certain intermediates. These computational predictions turner out to be consistent with experimental data obtained with transgenic plants ectopically overexpressing 3-hydroxy-3-methylglutary-coenzyme A reductase (*HMG1* gene). The opportunities and limitations of incorporating mathematical modeling into the design of approaches to engineer sterol biosynthesis are discussed.

## INTRODUCTION

Free sterols, sterol glycosides and acyl sterol glycosides are major lipid constituents of the plasma membrane (PM) in plants, accounting for > 40 mol-% of the total lipids within the PM of Arabidopsis (Uemura et al., 1995). Sterols are also enriched within detergent-resistant microdomains (DRMs) within the PM that are enriched in saturated fatty acids, sphingolipids and sterols, thus making them less fluid than the surrounding lipid bilayer (Georg et al., 2005; Takahashi et al., 2016). These regions serve as specific docking sites for proteins involved in cellulose synthesis (Bessueille et al., 2009), cold acclimation (Minami et al., 2015) and auxin transport (Willemsen et al., 2003). Mutation of sterol C-24 methyltransferase (SMT1) and experiments with methyl-β-cyclodextrin, a reagent that extracts sterols from membranes, have demonstrated that sterol depletion compromises the integrity of DRMs, thereby leading to the de-localization of DRM-associated proteins (Men et al., 2008; Kierszniowska et al., 2009). Sterols are also involved in several aspects of plant development, including male gametophyte and seed development (Jiang and Lin, 2013), embryo development, meristem function, hypocotyl elongation (Jang et al., 2000), stomatal development (Qian et al., 2013) and polar auxin transport (Willemsen et al., 2003; Men et al., 2008).

The biosynthesis of plant sterols has been the subject of a large number of studies, while the regulation of the pathway is much less understood. The mevalonate (MVA) pathway, which provides the bulk of precursors for the synthesis of sterols in plants, consists of eight enzymatic steps, yielding isopentenyl diphosphate (IPP) and dimethylallyl diphosphate (DMAPP) from acetyl-CoA (Hemmerlin et al., 2012) (Figure 1). One molecule of DMAPP and two molecules of IPP are then used to synthesize farnesyl diphosphate (FPP), which is further converted to squalene by squalene synthase (Kribii et al., 1997; Busquets et al, 2008). Squalene epoxidase generates squalene epoxide, which is the substrate for several triterpene synthases (Rasbery et al., 2007). The majority of squalene epoxide in leaves is usually converted to cycloartenol (> 85% of all carbon produced by the MVA pathway), in the first committed step of the core sterol pathway (Babichuk et al., 2008). In the major route toward sterol synthesis in plants, the side chain of cycloartenol is methylated in a reaction catalyzed by SMT1 (Shi et al., 1996; Bouvier-Navé et al., 1997; Diener et al., 2000; Nes et al., 2003), and the sterol nucleus is subsequently modified. A minor route for sterol synthesis modifies cycloartenol to give cholesterol (Figure 1). The enzymes involved in further reshaping the sterol nucleus use products of both the cholesterol and cycloartenol methylation pathways as substrates. The downstream reactions involve a sterol demethylase complex (Pascal et al., 1993; Rondet et al., 1999; Darnet et al, 2001; Darnet and Rahier, 2004; Rahier et al., 2006; d’Andréa et al., 2007; Li et al., 2007; Rahier, 2011), an isomerization (Rahier et al., 1989; Lovato et al., 2000), a second demethylation (Taton and Rahier, 1991a; Kushiro et al. 2001; Kim et al., 2005), a double bond reduction (Taton et al., 1989; Schrick et al., 2000; Jang et al., 2000), and a second isomerization (Grebenok et al., 2000; Souter et al., 2002). At the following branch point, 4α-methyl-5α-cholest-7-en-3β-ol and 24-methylenelophenol can be converted, via a series of reactions, to cholesterol and campesterol, respectively. Alternatively, 24-methylenelophenol can undergo a second methylation of the side chain to yield 24-ethylidenelophenol, catalyzed by a second sterol C-24 methyltransferase (SMT2) (Shi et al., 1996; Husselstein et al., 1996; Zhou et al., 2003). Enzymes involved in the late steps of sterol biosynthesis appear to operate on all three branches. These reactions involve a second sterol demethylase complex (Pascal et al., 1993; Rondet et al., 1999; Darnet et al, 2001; Darnet and Rahier, 2004; Rahier et al., 2006; d’Andréa et al., 2007; Li et al., 2007), a sterol desaturase (Gachotte et al., 1996; Taton and Rahier, 1996; Husselstein et al., 1999; Choe et al., 1999a), and two sterol reductases (Taton and Rahier, 1991b; Lecain et al., 1996; Klahre et al., 1998; Choe et al., 1999b; Choe et al., 2000), thereby leading to the major end products of the three branches, cholesterol, campesterol and β-sitosterol. Monooxygenases of the CYP710A family catalyze desaturations that generate the minor end products crinosterol/brassicasterol and stigmasterol (Morikawa et al., 2006; Arnqvist et al., 2008) (Figure 1).

**Figure 1.**
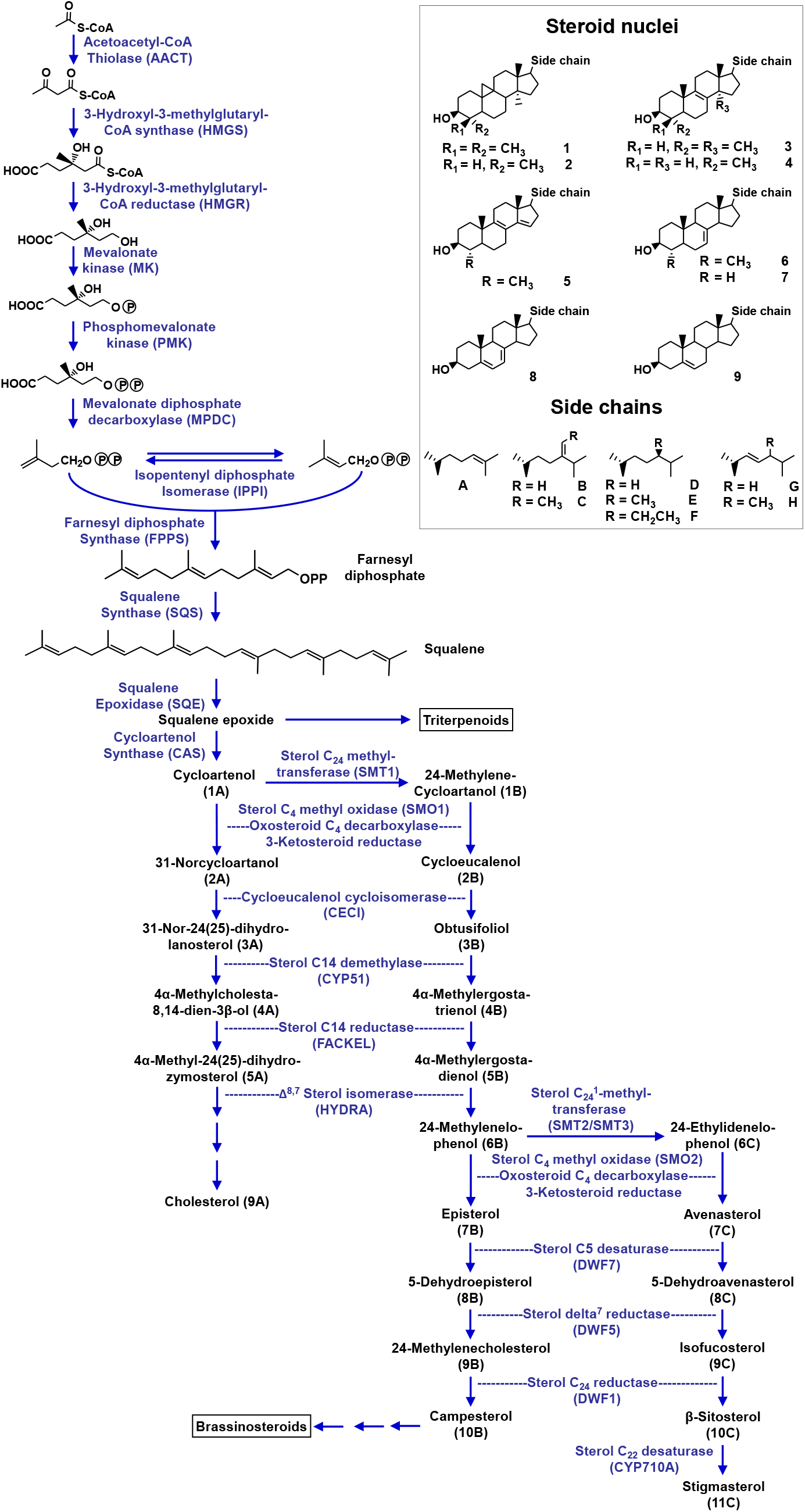
Outline of the core sterol biosynthetic pathway in plants.

Past studies on sterol accumulation have typically focused on a single organ or tissue cultured cells for analysis, but did not report on organ-specific compositional differences (Moreau et al., 2018). In the present study, we quantified sterols in four organs (root, stem, leaf, seed) of *Arabidopsis thaliana* (L.). Based on these data sets we developed kinetic mathematical models of sterol biosynthesis to capture flux distribution and pathway regulation in different organs. Simulations indicated that an increased flux through the sterol pathway would not only result in an increase of sterol end products but also a concomitant build-up of certain intermediates. These computational predictions were consistent with experimental data obtained with transgenic plants ectopically overexpressing 3-hydroxy-3-methylglutary-coenzyme A reductase (*HMG1* gene). The opportunities and limitations of incorporating mathematical modeling into the design of approaches to engineer sterol biosynthesis are discussed.

## RESULTS AND DISCUSSION

### Sterol profiles differ significantly across organs

Tissue samples were harvested at defined stages of Arabidopsis development (see Materials and Methods for details). In roots, the concentration of total sterols was 0.6 micromoles per gram fresh weight (μmol ·g FW^−1^) (Table 1). β-Sitosterol (49.6%), stigmasterol (22.5%) and campesterol (19.9%) were the major constituents, with cholesterol (1.3%), sterol pathway intermediates (total of 6.7 %; cycloartenol being the most abundant at 5.1%), and off-pathway triterpenoids (β-amyrin at 2.3% and lupeol at 0.4%) accumulating at lower levels (Table 1). Total sterols amounted to 0.4 μmol ·g FW^−1^ in rosette leaves (Table 1). The major constituents were β-sitosterol (72.8%) and campesterol (17.0%), but sterol pathway intermediates (isofucosterol at 5.3% and cycloartenol at 4.5%) were also detected with considerable abundance. The total sterol level in Arabidopsis stems was 0.99 μmol ·g FW^−1^. β-Sitosterol (66.9%), campesterol (14.7%) and brassicasterol (9.2%) were highly abundant, while cholesterol (2.2%), sterol pathway intermediates (cycloartenol at 6.5%), and off-pathway triterpenoids (β-amyrin at 5.3%) were minor contributors to the sterol profile. Arabidopsis seeds had the highest total sterol content of the investigated samples (8.2 μmol ·g FW^−1^) (Table 1). Of the sterol pathway end products, β-sitosterol (71.2%) and campesterol (16.1%) were major constituents, with cholesterol (3.4%), stigmasterol (1.3%) and brassicasterol (1.0%) being minor contributors to the profile. The amounts of sterol pathway intermediates (isofucosterol at 3.8% and cycloartenol at 2.6%) and off-pathway triterpenoids (β-amyrin at 1.9% and lupeol at 0.2%) were comparatively low (Table 1).

**Table 1.**
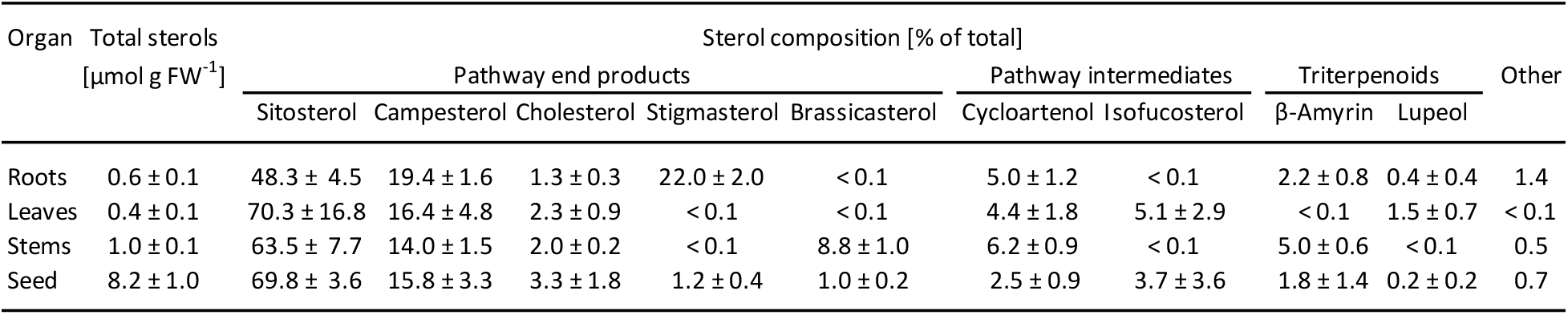
Sterol and triterpenoid levels in different Arabidopsis organs (n = 6-10; values indicate averages and standard errors).

| Organ | Total sterols<br>[μmol g FW<sup>-1</sup>] | Sterol composition [% of total] |  |  |  |  |  |  |  |  |  |
| --- | --- | --- | --- | --- | --- | --- | --- | --- | --- | --- | --- |
|  |  | Pathway end products |  |  |  |  | Pathway intermediates |  | Triterpenoids |  | Other |
|  |  | Sitosterol | Campesterol | Cholesterol | Stigmasterol | Brassicasterol | Cycloartenol | Isofucosterol | β-Amyrin | Lupeol |  |
| Roots | 0.6 ± 0.1 | 48.3 ± 4.5 | 19.4 ± 1.6 | 1.3 ± 0.3 | 22.0 ± 2.0 | <0.1 | 5.0 ± 1.2 | <0.1 | 2.2 ± 0.8 | 0.4 ± 0.4 | 1.4 |
| Leaves | 0.4 ± 0.1 | 70.3 ± 16.8 | 16.4 ± 4.8 | 2.3 ± 0.9 | <0.1 | <0.1 | 4.4 ± 1.8 | 5.1 ± 2.9 | <0.1 | 1.5 ± 0.7 | <0.1 |
| Stems | 1.0 ± 0.1 | 63.5 ± 7.7 | 14.0 ± 1.5 | 2.0 ± 0.2 | <0.1 | 8.8 ± 1.0 | 6.2 ± 0.9 | <0.1 | 5.0 ± 0.6 | <0.1 | 0.5 |
| Seed | 8.2 ± 1.0 | 69.8 ± 3.6 | 15.8 ± 3.3 | 3.3 ± 1.8 | 1.2 ± 0.4 | 1.0 ± 0.2 | 2.5 ± 0.9 | 3.7 ± 3.6 | 1.8 ± 1.4 | 0.2 ± 0.2 | 0.7 |

The information regarding sterol profiles in the literature is fragmentary, as most publications focus on a single organ. Roots were reported to contain high concentrations of β-sitosterol (70-80%), medium-high quantities of campesterol and stigmasterol (14-18 and 10-20%, respectively), and a relatively low cholesterol content (2-12%) (Wewer et al., 2011), which is generally consistent with our data, although the β-sitosterol content in our assays was significantly lower (50%). For stems and leaves, the reported composition is also in good agreement with our measurements (60-80% β-sitosterol, 14-18% campesterol, 0.5-2.0% cholesterol, and 0.5 - 1.0% stigmasterol) (Patterson et al., 1993; Gachotte et al., 1995; Schrick et al., 2000). High β-sitosterol amounts (75-80%) and medium-high quantities of campesterol (12-17%) were determined for seeds (stigmasterol and cholesterol levels not reported) (Chen et al., 2007), once again concurring with our data. The triterpenoids β-amyrin and lupeol were present in low quantities in most samples but, in stems, β-amyrin contributed more significantly (5% of total sterols and triterpenoids), which is consistent with other work on triterpenoids (Shan et al., 2008). To the best of our knowledge, the measurements presented here constitute the first data set directly comparing sterol and triterpenoid profiles across multiple organs, with the additional advantage of being obtained with a single analytical platform.

### Kinetic mathematical models of Arabidopsis sterol biosynthesis predict accumulation of pathway intermediates in transgenic plants with increased precursor availability

The pathway scheme shown in Figure 1 served as the conceptual foundation to develop proof-of-concept-level kinetic mathematical models of sterol biosynthesis in Arabidopsis. As a first step, the available information regarding kinetic constants and other biochemical knowledge (such as inhibition or activation) was assembled from the literature (Table 2). The Michaelis constant (*K*_M_) had been determined experimentally for many of the relevant enzymes but the turnover number (*k*_cat_) had to be estimated in many cases. For these approximate calculations, the reaction rate (*V*_max_) was taken into account when available (*k*_cat_ = *V*_max_ / [E]) (Table 2).

**Table 2.** Kinetic properties of sterol pathway enzymes (values with asterisks have been determined experimentally, all others are estimated). The units are uM (*K*_M_), nmol min^−1^ mg protein^−1^ (*V*_max_), and s^−1^ (*k*_cat_). The numbering of enzymes is in accordance with the scheme shown in Figure 1. Abbreviations: DMAPP, dimethylallyl diphosphate; FPP, farnesyl diphosphate; GPP, geranyl diphosphate; HMG-CoA, 3-hydroxy-3-methylglutaryl-coenzyme A; IPP, isopentenyl diphosphate; MP, microsomal preparation; MVA, mevalonate; PNE, purified native enzyme; PRE, purified recombinant enzyme.

| # | Enzyme | $K_M$ | $V_{\text{max}}$ | $k_{\text{cat}}$ | Comments | References |
| --- | --- | --- | --- | --- | --- | --- |
| 1 | Acetoacetyl-CoA thiolase (AACT) | 770* (Candida tropicalis; PNE) | - | 2.1* |  | Kanayama et al. (1997) |
| 2 | 3-Hydroxy-3-methylglutaryl-CoA synthase (HMGS) | 43* (Brassica juncea; PNE) | 470* | 0.415* | competitive inhibition by HMG-CoA ($K_i = 9$) | Nagegowda et al. (2004) |
| 3 | 3-Hydroxy-3-methylglutaryl-CoA reductase (HMGR) | 27* (Raphanus sativus; PNE) | 11* | 0.02 | competitive inhibition by MVA ($K_i = 990$) | Bach et al. (1986) |
| 4 | Mevalonate kinase (MK) | 76* (Catharanthus roseus; PNE) | 13.6* | 0.02 | competitive inhibition by FPP ($K_i = 0.1$) | Schulte et al. (2000) |
| 5 | Phosphomevalonate kinase (PMK) | 42* (Hevea brasiliensis; PNE) | 2.6* | 0.02 |  | Skilleter et al. (1971) |
| 6 | Mevalonate diphosphate decarboxylase (MPDC) | 10* (Cymbopogon citratus; PNE) | 1.12* | 0.02 |  | Lalitha et al. (1985) |
| 7 | Isopentenyl diphosphate: dimethylallyl diphosphate isomerase (IPPI) | 5.1* IPP, 17.0* DMAPP (Cinchona robusta; PNE) | 19.8* | 0.89* | uni-uni mechanism | Ramos-Valdivia et al. (1997) |
| 8 | Farnesyl diphosphate synthase (FPPS) | 15.3* IPP, 9.0* DMAPP, 1.8* GPP (Abies grandis; PNE) | 0.84* | 1.0* IPP, 9.0* DMAPP |  | Tholl et al. (2001) |
| 9 | Squalene synthase (SQS) | 9.5* FPP (Nicotiana tabacum; PNE) | - | 0.53* |  | Hanley and Chappell (1992) |
| 10 | Squalene monooxygenase / squalene epoxidase (SQE) | 7.7* (Homo sapiens; PRE) | - | 0.0183* |  | Laden et al. (2000) |
| 11 | Cycloartenol synthase (CAS) | 125* (Zea mays; PNE) | - | 0.015 |  | Abe et al. (1988) |
| 12 | Cycloartenol C24 methyltransferase (SMT1) | 30* (Zea mays; PNE) | 0.02* | 0.01* | competitive inhibition by 24-methylenecycloartenol ($K_i =$ competitive inhibition by 24-ethylidenelophenol ($K_i = 50$) competitive inhibition by sitosterol ($K_i = 100$) | Nes et al. (2003); Zhou and Nes (2003); Neelandankan et al. (2009); Parker and Nes (1992) |
| 13 | Sterol 4α demethylase complex (SMO1) | 500* (Zea mays; MP) | 0.135* | 0.06 | competitive inhibition by sitosterol ($K_i = 100$) | Pascal et al. (1993); Rahier et al. (2006); d'Andréa et al. (2007) |
| 14 | Cycloeucalenol cycloisomerase (CECI) | 100* (Zea mays; MP) | 0.6* | 0.05 |  | Rahier et al. (1989) |
| 15 | Obtusifoliol C14 demethylase (CYP51) | 160* (Zea mays; MP) | 0.065* | 0.05 |  | Taton and Rahier (1991) |
| 16 | Sterol C14 reductase (FACKEL) | 100* (Zea mays; MP) | 0.23* | 0.05 |  | Taton et al. (1989) |
| 17 | C-8,7 Sterol isomerase (HYDRA) | 75* (Zea mays; MP) | 0.2* | 0.05 |  | Rahier et al. (1997) |
| 18 | Sterol C24-methyltransferase (SMT2) | 28* methylene lophenol (Arabidopsis thaliana; PRE) | - | 0.01* | competitive inhibition by sitosterol ($K_i = 300$) | Zhou and Nes (2003); Parker and Nes |
| 19 | $\Delta^7$-Sterol C5-desaturase (DWF7) (campesterol) | 115* Cholesten-7-en-3β-ol (Zea mays; MP) | 0.13 | 0.06 | | Taton and Rahier (1996) |
| 20 | Sterol $\Delta^7$-reductase (DWF5) (campesterol branch) | 460* (Zea mays; MP) | 0.61 | 0.06 | | Taton and Rahier (1991) |
| 21 | Sterol C24-reductase (DWF1) (campesterol branch) | 150 | - | 0.06 |  | - |
| 22 | Conversion of campesterol into brassinosteroids | (considered as not carrying significant flux) | - | - |  | - |
| 23 | Sterol 4α demethylase complex (SMO2) | 480* (Zea mays; MP) | 0.16 | 0.06 | | Pascal et al. (1993); Pascal et al. (1994) |
| 24 | $\Delta^7$-Sterol C5-desaturase (DWF7) (sitosterol branch) | 140* Stigmasta-7,24(24')-dien-3β-ol (Zea mays; MP) | 0.13 | 0.06 | | Taton and Rahier (1996) |
| 25 | Sterol $\Delta^7$-reductase (DWF5) (sitosterol branch) | 460* (Zea mays; MP) | 0.61 | 0.06 | | Taton and Rahier (1991) |
| 26 | Sterol C24-reductase (DWF1) (sitosterol branch) | 150 | - | 0.0018 |  | - |
| 27 | Sterol C22-desaturase (CYP710A) | 1.0* (Arabidopsis thaliana; PRE) | - | 0.0001 |  | Morikawa et al. (2006) |
| 28 | Sterol C24-methyltransferase (SMT2) | 28 episterol (Arabidopsis thaliana; PRE) | - | 0.01* |  | Nes et al. (2003) |
| 29 | Sterol C24-methyltransferase (SMT2) | 28 5-dehydroepisterol (Arabidopsis thaliana; PRE) | - | 0.01* |  | Nes et al. (2003) |
| 30 | Sterol C24-methyltransferase (SMT2) | 28 24-methylene cholesterol (Arabidopsis thaliana; PRE) | - | 0.01* |  | Nes et al. (2003) |

As a second step, rate equations were generated based on the Michaelis-Menten kinetics. For example, the change in the pool size of squalene (sterol biosynthesis intermediate) is determined by the rates of formation (v_1_) and turnover (v_2_) and can be expressed as follows:

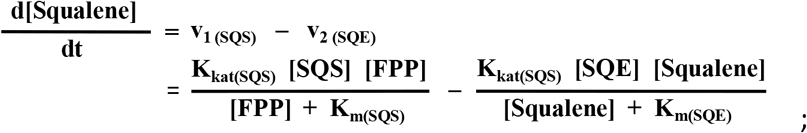

with FPP, farnesyl diphosphate; SQE, squalene epoxidase; SQS, squalene synthase.

This formalism was used for the description of a total of 27 enzymatic reactions, with appropriate equations for unidirectional and reversible reactions, multi-substrate reactions (e.g., ternary complex or ping-pong mechanisms), and, if known, feedback inhibition. Numerical solutions for the resulting system of ordinary differential equations (ODEs) were calculated by an iterative process of approximation and error correction (MATLAB software package, MathWorks).

As a third step, the concentration of each enzyme in each of the five organs of interest had to be estimated. The ideal - but unfortunately too costly - approach would have been organ-specific quantitative proteomics data. We decided to infer enzyme concentrations from gene expression data instead, which, based on our previous work (Rios-Estepa et al., 2010; Lange and Rios-Estepa, 2014), can be employed as a first approximation. The organ-specific gene expression data sets employed in the current work had been made available as part of a prior publication (Schmid et al., 2005). The factor to convert gene expression levels (unitless) to enzyme concentrations (in μM) was determined by evaluating computationally which combination of values predicted sterol concentrations most closely matching those determined experimentally (a conversion factor of 6,800 was chosen) (Table 3).

**Table 3.**
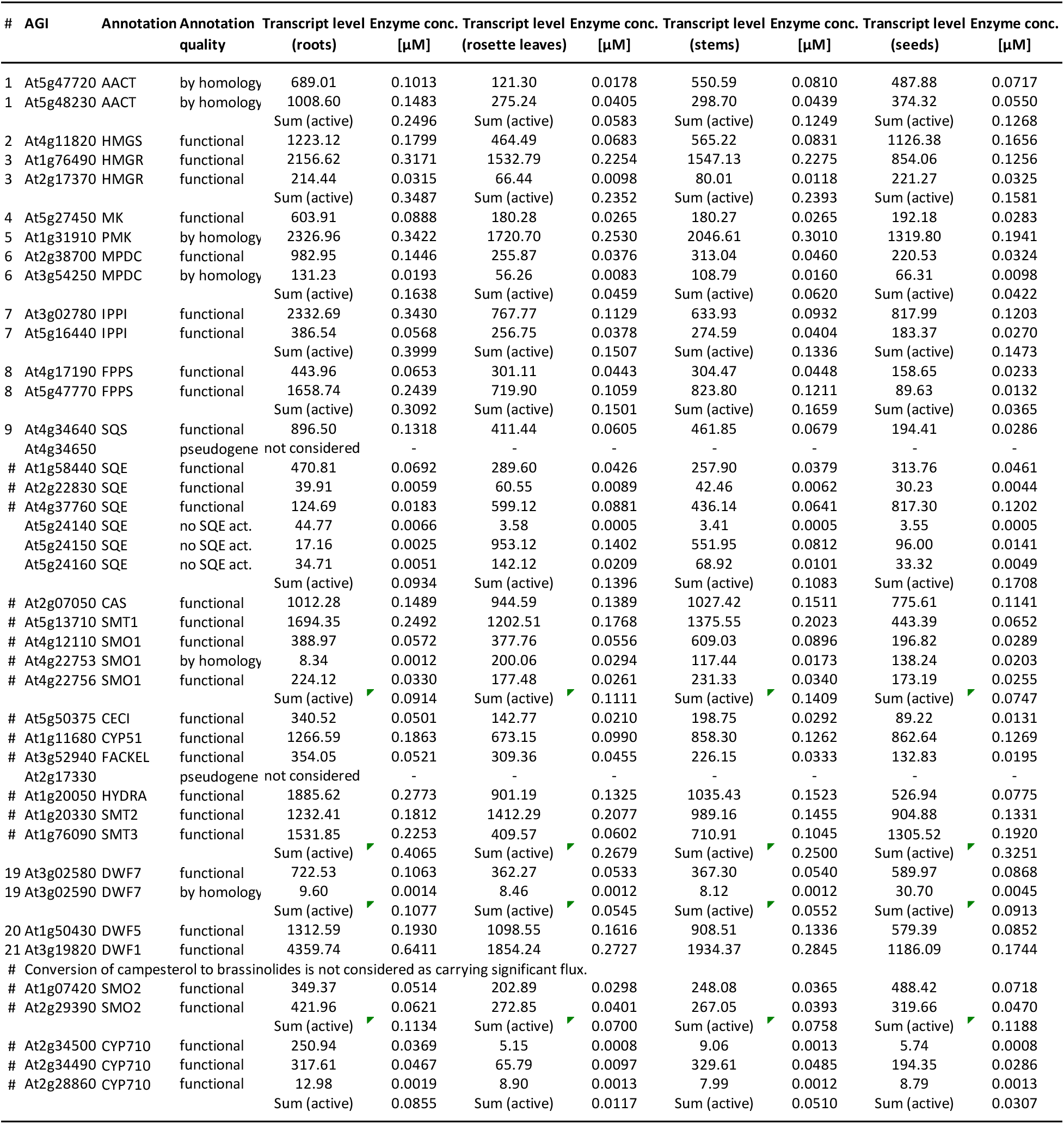
Expression levels of sterol biosynthetic genes in different organs and conversion into enzyme concentrations (for legend of abbreviations see Table 2). The numbering of genes is in accordance with that of the corresponding enzymes in Figure 1.

As a fourth step, model adjustments were made to reflect an additional consideration. A factor (termed Kc40) was introduced to account for the esterification of sterol pathway end products (β-sitosterol, stigmasterol, campesterol and cholesterol) and/or incorporation into biological membranes, which reduces the concentration of free sterols and thereby the potential for acting as feedback inhibitors of other biosynthetic enzymes (in particular sterol methyltransferases). Simulations were performed assuming a 50 d (7 week) growth period. The full details of the computational approaches used in this study are available in Supplemental Methods and Data File S1).

Our model for root sterol accumulation predicted high β-sitosterol levels (0.29 μmol · g FW^−1^) and relatively low concentrations for campesterol and stigmasterol (0.05 and 0.02 μmol ·g FW^−1^) (Figure 2A). These values are consistent with experimental data for β-sitosterol (0.29 μmol ·g FW^−1^) but too low for campesterol and stigmasterol (experimental: 0.12 and 0.13 μmol ·g FW^−1^, respectively). The leaf model also predicted β-sitosterol to accumulate as the most prominent sterol (0.30 μmol ·g FW^−1^), campesterol at low levels (0.09 μmol ·g FW^−1^), and all other sterols at very low concentrations (< 0.05 μmol ·g FW^−1^) (Figure 2B). This was consistent with experimental data (β-sitosterol and campesterol at 0.30 and 0.07 μmol ·g FW^−1^, respectively). The predictions for stems were high β-sitosterol levels (0.65 μmol ·g FW^−1^) (very close to the measured concentration of 0.66 μmol ·g FW^−1^), relatively high campesterol quantities (0.19 μmol ·g FW^−1^) (close to the measured concentration of 0.15 μmol ·g FW^−1^), while other sterols were predicted to play less prominent roles (also in line with experimental data) (Figure 2C). Our model for sterol biosynthesis in seeds predicted high concentrations of β-sitosterol (5.8 μmol ·g FW^−1^) and considerable quantities of campesterol (1.3 μmol ·g FW^−1^) and stigmasterol (0.4 μmol ·g FW^−1^) as well, which matches experimental values for β-sitosterol and campesterol well (5.84 and 1.33 μmol ·g FW^−1^, respectively), but is too high for stigmasterol (experimental: 0.10 μmol ·g FW^−1^) (Figure 2D). Overall, the predicted sterol profiles were a fairly good match of the experimentally determined sterol accumulation patterns across organs. This is quite remarkable when considering that our simulations use numerous assumptions, the quality of which can be substantially improved in future work. For example, the enzyme concentrations were estimated based on previously published gene expression patterns (Schmid et al., 2005), and, while we attempted to use similar growth conditions as reported in this publication, it is likely that the crude gene-to-protein concentration conversion lacked accuracy. It is also conceivable that the concentrations of sterol end products that directly affect biosynthetic enzymes by feedback inhibition might be estimated incorrectly. Overall, we feel that our proof-of-concept models certainly have utility for simulating sterol profiles and are an excellent first step toward truly predictive kinetic models of sterol biosynthesis.

**Figure 2.**
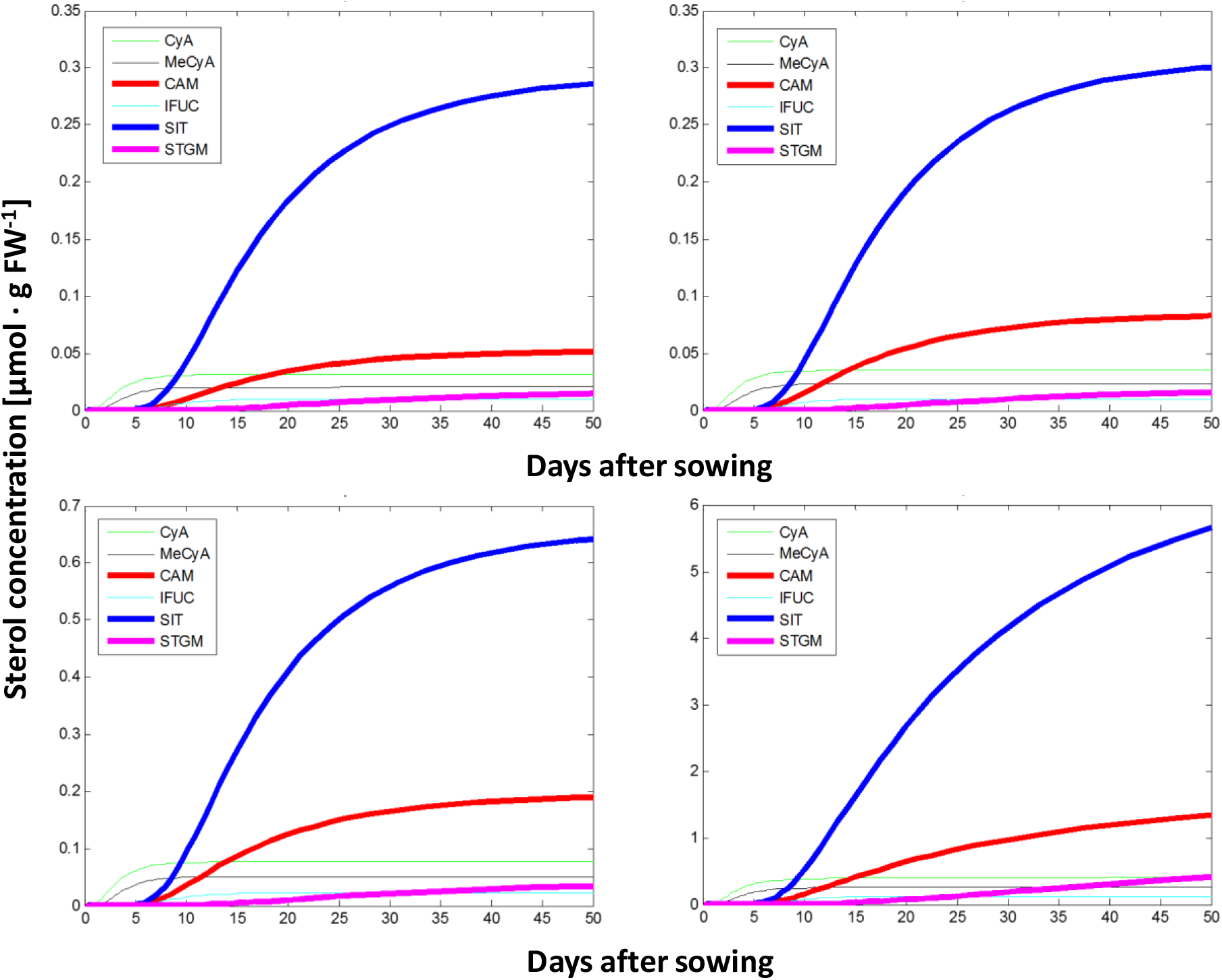
Prediction of sterol profiles, using kinetic mathematical modeling, in different organs of Arabidopsis. **A**, roots; **B**, rosette leaves; **C**, stems; and **D**, seeds. Abbreviations: CAM, campesterol; CyA, cycloartenol; IFUC, isofucosterol; MeCyA, 24-methylenecycloartanol; SIT, β-sitosterol; STGM, stigmasterol.

### Ectopic HMG1 overexpression leads to increases in sterol levels but also to the accumulation of certain pathway intermediates

Building on the initial successes with developing organ-specific models of sterol biosynthesis, we attempted to predict the effects of gene overexpression on sterol profiles in transgenic Arabidopsis plants. We recently reported on the generation and analysis of transgenics with varying levels of expression of genes involved in isoprenoid precursor biosynthesis (Lange et al., 2015) and selected the HMG1-6.1.7 line, which was shown to overexpress the *HMG1* gene in rosette leaves, for further analysis (no other organs had been analyzed before). For the current study, we employed quantitative real-time PCR to assess *HMG1* transcript levels. The *HMG1* gene was found to be overexpressed in roots, rosette leaves, stems and seeds (25-, 26-, 49-, and 10-fold, respectively) (Figure 3A). Our prior work with kinetic models indicated that the amplitude of transcript level variation appears to be larger than that of enzyme concentrations in comparisons of wild-type versus transgenic plants (Rios-Estepa et al., 2008; Rios-Estepa et al, 2010). We therefore divided the fold-change for *HMG1* overexpression by a factor of 5 to estimate the change in the concentration of the corresponding protein (5.0-, 5.2-, 9.8-, and 2.0-fold for roots, leaves, stems and seeds, respectively). The only other adjusted model parameters were kc40 (removes free sterols and decreasing feedback inhibition of biosynthetic enzymes by accumulating end products) and the initial amount of acetyl-coenzyme A precursor available for sterol pathway enzymes to act upon (reflects the experimentally determined increases of sterol content in *HMG1* overexpressors over wild-type controls; 3.7-fold in roots, 3.0-fold in rosette leaves, 3.3-fold in stems, and 1.6-fold in seeds) (Figure 3B). A detailed description of model parameters is given in Supplemental Methods and Data File S1).

**Figure 3.**
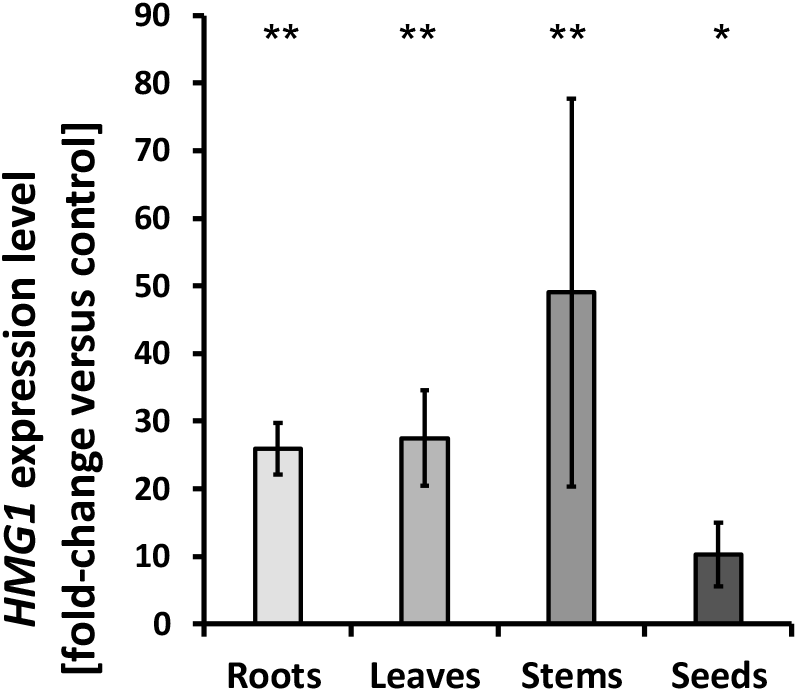
Analysis of Arabidopsis organ samples of wild-type controls and plants overexpressing the *HMG1* gene. **A**, Expression patterns of the *HMG1* gene (n = 3) and **B**, sterol content (n = 10) (both expressed as fold-change overexpressor versus control). Standard errors shown as bars; asterisks indicate the level of significance in a Student’s t test (*, P ≤ 0.01; and **, P ≤ 0.001).

Root sterol concentrations for *HMG1* overexpressors were predicted to amount to 0.98 μmolg FW^−1^ for β-sitosterol and 0.18 μmol ·g FW^−1^ for campesterol (Figure 4A), which were also the principal sterols in experimental samples (0.99 and 0.40 μmol ·g FW^−1^, respectively). Our rosette leaf model predicted an increase (when compared to wild-type controls) of β-sitosterol and campesterol (0.65 and 0.18 μmol ·g FW^−1^, respectively) (Figure 4B), which was also close to experimentally determined values (0.64 and 0.14 μmol ·g FW^−1^, respectively). When simulating stem sterol profiles, β-sitosterol and campesterol concentrations were predicted to amount to 1.64 and 0.49 μmol ·g FW^−1^, respectively (Figure 4C), which closely matched experimental data (1.64 and 0.40 μmol ·g FW^−1^, respectively). The model predictions for seeds were dramatically increased concentrations of β-sitosterol and campesterol (8.2 and 1.8 μmol ·g FW^−1^, respectively) (Figure 4D), again an excellent reflection of experimental measurements (8.31 and 1.88 μmol ·g FW^−1^, respectively). Interestingly, all organ-specific models predicted a significant increase in the proportion of the sterol pathway intermediates, in particular cycloartenol, 24-methylene-cycloartanol and isofucosterol, in *HMG1* plants compared to wild-type controls. While these intermediates account for only a relatively small percentage of the total sterols in wild-type plants, we measured significant increases in *HMG1* plants, compared to controls, in all organs (cycloartenol, 6-11-fold; 24-methylenecycloartanol, 5-20-fold; isofucosterol, 2.6-180-fold) (Figure 5A-C). The accumulation of intermediates of sterol biosynthesis has been reported repeatedly for transgenic plants with increased flux into the pathway (by overexpression of one or more genes involved in precursor supply) (Chappell et al., 1995; Holmberg et al., 2003; Lange et al., 2015). When a gene that encodes a protein that turns over an accumulated intermediate (such as SMO1 for cycloartenol and 24-methylenecycloartanol, SMT1 for cycloartenol, or DWF1 for isofucosterol) was overexpressed in plants already overexpressing *HMG1*, flux constraints could be partially removed and additional increases in sterol end products were observed (Holmberg et al., 2003; Lange et al., 2015). In summary, our models have been remarkably accurate with predicting sterol profiles across organs and genotypes. If more accurate data sets of enzyme concentrations were to be integrated into these models, the predictive power would likely increase as well. In our opinion, the fact that the accumulation of sterol pathway intermediates was correctly predicted indicates that kinetic modeling can point to potential flux bottlenecks, thereby suggesting experimental approaches toward flux enhancement. It will now be interesting to evaluate if a combination of modeling and experimentation has the potential to speed up metabolic engineering by being able to focus on the most promising avenues for the accumulation of desirable metabolites.

**Figure 4.**
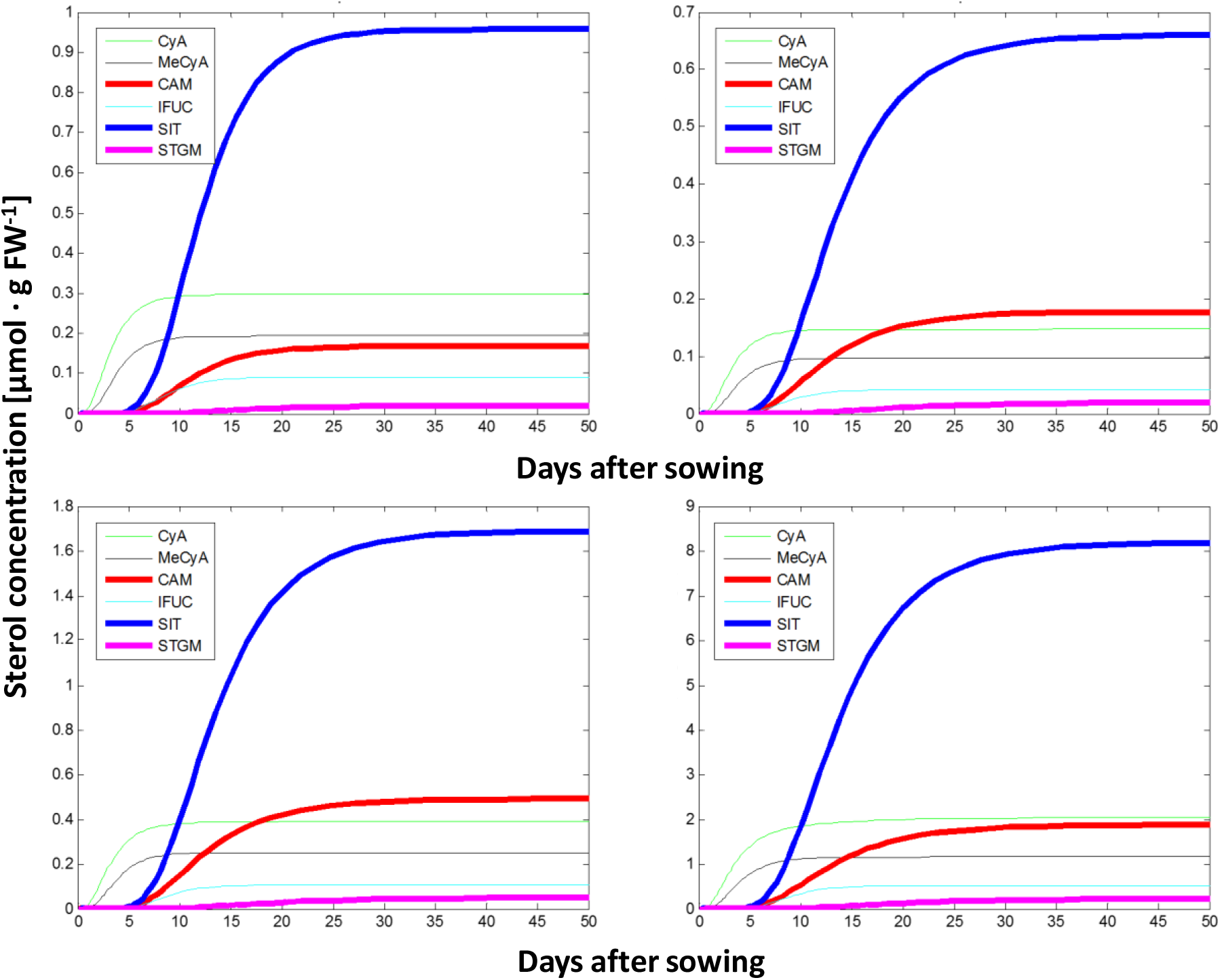
Prediction of sterol profiles, using kinetic mathematical modeling, in different organs of transgenic Arabidopsis plants overexpressing the *HMG1* gene. **A**, roots; **B**, rosette leaves; **C**, stems; and **D**, seeds. Abbreviations: CAM, campesterol; CyA, cycloartenol; IFUC, isofucosterol; MeCyA, 24-methylenecycloartanol; SIT, β-sitosterol; STGM, stigmasterol.

**Figure 5.**
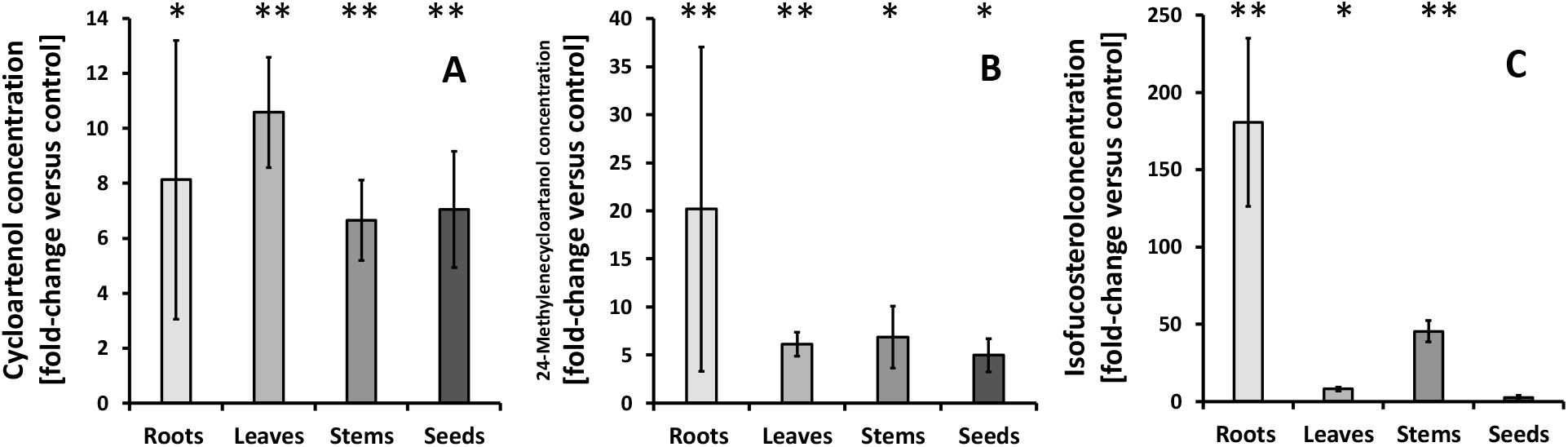
Analysis of sterol pathway intermediates of wild-type controls and plants overexpressing the *HMG1* gene. **A**, cycloartenol; **B**, 24-methylenecycloartanol; **C**, isofucosterol (all expressed as fold-change overexpressor versus control; n = 10). Standard errors shown as bars; asterisks indicate the level of significance in a Student’s t test (*, P ≤ 0.01; and **, P ≤ 0.001; n = 6-10).

## Materials and Methods

### Plant growth for harvest of aboveground tissue samples

Seeds of wild-type and homozygous transgenic lines were germinated in soil (6 × 6 cm pots) and maintained in a growth chamber (16 h day/8 h night photoperiod; 100 μmol m^−2^ s^−1^ light intensity at soil level; constant temperature at 23°C; 70% relative humidity). Pots were arranged in a random grid and rotated once a day. Rosette leaves were collected at growth stage 5.10 (just as the inflorescence starts to appear; Boyes et al., 2001); stems and siliques were collected at growth stage 6.50 from a separate set of plants. Developing seeds were separated from siliques using a rapid separation procedure (Bates et al., 2013). Mature seeds were collected at growth stage 9.70 from yet another set of plants. Leaf and stem tissue samples were used for both qPCR and metabolite analyses. Immature seeds were used for qPCR expression analysis (during seed filling), whereas mature seeds were used for metabolite analyses. Samples were shock-frozen in liquid nitrogen immediately following harvest, the frozen tissue samples homogenized with mortar and pestle in the presence of liquid nitrogen, and aliquots of the resulting homogenate stored in 2 ml Eppendorf tubes at −80°C. Samples were further homogenized (Ball Mill MM301, Retsch) before subsequent analyses. Samples were harvested from three independent experiments (biological replicates).

### Plant growth for harvest of root tissue

Plants were grown in an aeroponic culture system (according to Vaughan et al., 2011) using the same lighting conditions as described above. Briefly, plants were grown in soil for 3 weeks, then removed from soil and transplanted into 50 ml conical vials containing Seramis clay granulate, with holes punched into the bottom for water access. The vials were placed in a solution of half-strength MS basal salt for 10 min once every 48 h. Pots and vials were arranged in a random grid and rotated once a day. After 3 weeks of growth in the aeroponic system, plants were removed from the vials and roots were separated from the Seramis granules for harvest. Root tissue was harvested, homogenized and stored as described above.

### Quantitative real-time PCR

Root, rosette and stem RNA was extracted using the Trizol reagent (Life Technologies) according to the manufacturer’s instructions. RNA extraction from seeds involved a two-step Trizol-based procedure (Meng and Feldmann, 2010). Isolated RNA (1,000 ng) was treated with RNase-free DNase (Thermo Scientific) and first strand cDNA synthesized using Superscript III reverse transcriptase (Life Technologies). In a 10-μL quantitative PCR reaction, concentrations were adjusted to 150 nM (primers), 1 × Power SYBR Green PCR Master Mix (Life Technologies), and 10 x diluted first strand cDNA as template (primer sequences provided in Table S1). Reactions were performed in a 96-well optical plate at 95°C for 10 min, followed by 40 cycles of 95°C for 15 s and 60°C for 10 min in a 7500 Real-Time PCR system (Life Technologies). Fluorescence intensities of three independent measurements (technical replicates) were normalized against the ROX reference dye (ThermoFisher Scientific). Relative transcript levels were calculated based on the comparative CT method as specified in the manufacturer’s instructions (Life Technologies). β-Actin (At3g18780) served as the constitutively expressed endogenous control and the expression level of the corresponding wild-type allele was used as the calibrator in these calculations.

### Sterol analyses

Tissue homogenate (60–70 mg for root, leaf and stem; 10 mg for seed) was extracted at 75°C for 60 min with 4ml of CHCl_3_/MeOH (2:1, v:v; containing 1.25 mg ·L^−1^ epi-cholesterol as internal standard). Samples were evaporated to dryness (EZ2-Bio, GeneVac), and the remaining residue was saponified at 90°C for 60 min in 2 ml of 6% (w/v) KOH in MeOH. Upon cooling to room temperature, 1 ml of hexane and 1 ml of H_2_O were added, and the mixture was shaken vigorously for 20 s. Following centrifugation (3,000 × *g* for 2 min) for phase separation, the hexane phase was transferred to a 2 ml glass vial, and the aqueous phase re-extracted with 1 ml of hexane as above. The mixture was centrifuged as above, the hexane phase removed and added to the 2 ml glass vial containing the hexane phase from the first extraction. The combined organic phases were evaporated to dryness as above, 50 μl of N-Methyl-N-(trimethylsilyl)trifluoroacetamide were added to the residue, the mixture was shaken vigorously for 20 s, and the sample then transferred to a 2 ml glass vial with a 100-μl conical glass insert. After capping the vial, the reaction mixture was incubated at room temperature for 5 min. Gas chromatography-mass spectrometry analyses were performed on an Agilent 6890N gas chromatograph coupled to an Agilent 5973 inert mass selective detector (MSD) detector. Samples were loaded (injection volume 1 μl) with a LEAP CombiPAL autosampler onto an Agilent HP-5MS fused silica column (30 m x 250 mm; 0.25-mm film thickness). The temperatures of the injector and MSD interface were both set to 280°C. Analytes were separated at a flow rate of 1 ml ·min^−1^, with He as the carrier gas, using a thermal gradient starting at 170°C (hold for 1.5 min), a first ramp from 170°C to 280°C at 37°C/min, a second ramp from 280°C to 300°C at 1.5°C/min, and a final hold at 300°C for 5.0 min. Eluting metabolites were fragmented in electron impact mode with an ionization voltage of 70 eV and data was acquired using MSD ChemStation software (revision D.01.02.SP1, Agilent Technologies). Analytes were identified based on their mass fragmentation patterns by comparison with those of authentic standards using the National Institute of Standards and Technology Mass Spectral Search Program (version D.05.00). Peak areas were obtained from the total ion chromatogram for all detectable peaks with a sterol mass fragmentation signature. Raw data were exported to Microsoft Excel and peak areas normalized to tissue mass and internal standard. A blank injection was performed after each sample run, and the background signal from the blank subtracted from the sample values for the entire run. Prior to sample analyses, and then after every 20 samples, a standard mix was run to evaluate the reproducibility of the analyses. Absolute quantitation of sterols was achieved based on calibration curves obtained with authentic standards.

#### Development of organ-specific kinetic models of sterol biosynthesis

Publicly available information regarding the biochemical properties of enzymes involved in sterol biosynthesis (Michaelis constant, turnover number, and inhibition constant) was assembled from the literature and the BRENDA repository (https://www.brenda-enzymes.org/). The information was then further consolidated by prioritizing kinetic values from Arabidopsis, Brassicaceae, plants and other eukaryotes (in this order). For each reaction of the sterol pathway, the concentration change of reactants over time was defined according to the Michaelis-Menten formalism, as developed by Briggs and Haldane (Fersht, 1985), and converted into ODEs. These were then integrated into a system of interdependent ODEs and numerical solutions obtained in MATLAB version 7.12.0.635 (MathWorks) using the ode15s solver. The full MATLAB code, with explanations, is provided in Supplemental Methods and Data File S1. Simulations were performed assuming a 50 d (7 week) growth period. The following files, functions, parameters and variables were generated within MATLAB:

##### Script File

A set of commands that includes the vector for pathway metabolites, time span, and the vector of initial conditions. It calls the function (m-file) that solves the ODEs and produces the graphical outputs (monoterpene profiles).

##### Function (m-file)

Inputs: independent variable ***t*** (time span); vector of dependent variables ***x*** ([Metabolites]). Solves the set of ODEs with the initial values given in the vector of initial conditions. Returns the values of the independent variable in the vector ***t*** (time span) and the values of the dependent variables in the vector ***x*** ([Metabolites]). The vector of independent variables ***t*** is not equally spaced because the function (m-file) controls the step size.

##### Fixed parameters

Kinetic constants of enzymes involved in sterol biosynthesis.

##### Non-constant parameters (variables)

Enzyme concentrations for each organ and genotype. Factor kc40 to account for the conversion of frees sterols into sterol esters and incorporation into membranes, thereby reducing the concentration of sterol pathway end products that can act on biosynthetic enzymes via feedback inhibition. Initial amount of starting material (acetyl-CoA) in the script file (all other metabolite concentrations are set to 0 at t_0_).

## Acknowledgements

This work was supported by the National Science Foundation (grant no. NSF-MCB-090758 to B.M.L.).

## Author Contributions

BML conceived of the project. BCP and IL designed and performed the experimental work. RRE and RS developed the kinetic mathematical models. All authors were involved in data analysis. BML wrote the manuscript, with contributions from all other authors.

## REFERENCES

Arnqvist L, Persson M, Jonsson L, Dutta PC, Sitbon F (2008) Overexpression of CYP710A1 and CYP710A4 in transgenic *Arabidopsis* plants increases the level of stigmasterol at the expense of sitosterol. Planta 227: 309–317

Babiychuk E, Bouvier-Navé P, Compagnon V, Suzuki M, Muranaka T, Van Montagu M, Kushnir S, Schaller H (2008) Allelic mutant series reveal distinct functions for *Arabidopsis* cycloartenol synthase 1 in cell viability and plastid biogenesis. Proc Natl Acad Sci USA 105: 3163–3168

Bates PD, Jewell JB and Browse J (2013) Rapid separation of developing Arabidopsis seeds from siliques for RNA or metabolite analysis. Plant Methods 9: 9

Bessueille L, Sindt N, Guichardant M, Djerbi S, Teeri TT, Bulone V (2009) Plasma membrane microdomains from hybrid aspen cells are involved in cell wall polysaccharide biosynthesis. Biochem J 420: 93–103

Bouvier-Navé P, Husselstein T, Desprez T, Benveniste P (1997) Identification of cDNAs encoding sterol methyl-transferases involved in the second methylation step of plant sterol biosynthesis. Eur J Biochem 246: 518–529

Boyes DC, Zayed AM, Ascenzi R, McCaskill AJ, Hoffman NE, Davis KR, Görlach J (2001) Growth stage-based phenotypic analysis of Arabidopsis: a model for high throughput functional genomics in plants. Plant Cell 13: 1499–1510

Busquets A, Keim V, Closa M, Del Arco A, Boronat A, Arró M, Ferrer A (2008) *Arabidopsis thaliana* contains a single gene encoding squalene synthase. Plant Mol Biol 67: 25–36.

Chappell J, Wolf F, Proulx J, Cuellar R, Saunders C (1995) Is the reaction catalyzed by 3-hydroxy-3-methylglutaryl coenzyme A reductase a rate-limiting step for isoprenoid biosynthesis in plants? Plant Physiol 109: 1337–1343

Chen Q, Steinhauer L, Hammerlindl J, Keller W, Zou J (2007) Biosynthesis of phytosterol esters: identification of a sterol o-acyltransferase in Arabidopsis. Plant Physiol 145: 974–984

Choe S, Noguchi T, Fujioka S, Takatsuto S, Tissier CP, Gregory BD, Ross AS, Tanaka A, Yoshida S, Tax FE, Feldmann KA (1999a) The *Arabidopsis* dwf7/ste1 mutant is defective in the delta7 sterol C-5 desaturation step leading to brassinosteroid biosynthesis. Plant Cell 11: 207–221

Choe S, Dilkes BP, Gregory BD, Ross AS, Yuan H, Noguchi T, Fujioka S, Takatsuto S, Tanaka A, Yoshida S, et al (1999b) The Arabidopsis dwarf1 mutant is defective in the conversion of 24-methylenecholesterol to campesterol in brassinosteroid biosynthesis. Plant Physiol 119: 897–907

Choe S, Tanaka A, Noguchi T, Fujioka S, Takatsuto S, Ross AS, Tax FE, Yoshida S, Feldmann KA (2000) Lesions in the sterol delta reductase gene of *Arabidopsis* cause dwarfism due to a block in brassinosteroid biosynthesis. Plant J 21: 431–443

d’Andréa S, Canonge M, Beopoulos A, Jolivet P, Hartmann MA, Miquel M, Lepiniec L, Chardot T (2007) At5g50600 encodes a member of the short-chain dehydrogenase reductase superfamily with 11beta-and 17beta-hydroxysteroid dehydrogenase activities associated with Arabidopsis thaliana seed oil bodies. Biochimie 89: 222–229

Darnet S, Bard M, Rahier A (2001) Functional identification of sterol-4alpha-methyl oxidase cDNAs from *Arabidopsis thaliana* by complementation of a yeast erg25 mutant lacking sterol-4alpha-methyl oxidation. FEBS Lett 508: 39–43

Darnet S, Rahier A (2004) Plant sterol biosynthesis: identification of two distinct families of sterol 4alpha-methyl oxidases. Biochem J 378: 889–898

Diener AC, Li H, Zhou W, Whoriskey WJ, Nes WD, Fink GR (2000) Sterol methyltransferase 1 controls the level of cholesterol in plants. Plant Cell 12: 853–870

Fersht A (1985) Enzyme structure and mechanism, W.H. Freeman, New York, 475 pages

Gachotte D, Meens R, Benveniste P (1995) An Arabidopsis mutant deficient in sterol biosynthesis: heterologous complementation by ERG 3 encoding a delta7-sterol-C-5-desaturase from yeast. Plant J 8: 407–416

Gachotte D, Husselstein T, Bard M, Lacroute F, Benveniste P (1996) Isolation and characterization of an *Arabidopsis thaliana* cDNA encoding a delta 7-sterol-C-5-desaturase by functional complementation of a defective yeast mutant. Plant J 9: 391–398

Georg HB, Sherrier DJ, Weimar T, Louise VM, Hawkins ND, MacAskill A, Napier JA, Beale MH, Lilley KS, Dupree P (2005) Analysis of detergent-resistant membranes in Arabidopsis; evidence for plasma membrane lipid rafts. Plant Physiol 137: 104–116

Grebenok RJ, Ohnmeiss TE, Yamamoto A, Huntley ED, Galbraith DW, DellaPenna D (2000) Isolation and characterization of an *Arabidopsis thaliana* C-8,7 sterol isomerase: functional and structural similarities to mammalian C-8,7 sterol isomerase /emopamil-binding protein. Plant Mol Biol 38: 807–815

Hemmerlin A, Harwood JL, Bach TJ (2012) A raison d’être for two distinct pathways in the early steps of plant isoprenoid biosynthesis? Prog Lipid Res 51: 95–148

Hey SJ, Powers SJ, Beale MH, Hawkins ND, Ward JL, Halford NG (2006) Enhanced seed sterol accumulation through expression of a modified HMG-CoA reductase. Plant Biotechnol J 4: 219–229

Holmberg N, Harker M, Wallace AD, Clayton JC, Gibbard CL, Safford R (2003) Co-expression of N-terminal truncated 3-hydroxy-3-methylglutaryl CoA reductase and C24-sterol methyltransferase type 1 in transgenic tobacco enhances carbon flux towards end-product sterols. Plant J 36: 12–20

Husselstein T, Gachotte D, Desprez T, Bard M, Benveniste P (1996) Transformation of *Saccharomyces cerevisiae* with a cDNA encoding a sterol C-methyltransferase from Arabidopsis thaliana results in the synthesis of 24-ethyl sterols. FEBS Lett 381: 87–92

Husselstein T, Schaller H, Gachotte D, Benveniste P (1999) Delta7-sterol-C5-desaturase: molecular characterization and functional expression of wild-type and mutant alleles. Plant Mol Biol 39: 891–906

Jang JC, Fujioka S, Tasaka M, Seto H, Takatsuto S, Ishii A, Aida M, Yoshida S, Sheen J (2000) A critical role of sterols in embryonic patterning and meristem programming revealed by the fackel mutants of Arabidopsis thaliana. Genes Dev 14: 1485–1497

Jiang W and Lin W (2013) Brassinosteroid functions in Arabidopsis seed development. Plant Signal Behav 8: 10

Kierszniowska S, Seiwert B, Schulze WX (2009) Definition of Arabidopsis sterol-rich membrane microdomains by differential treatment with methyl-beta-cyclodextrin and quantitative proteomics. Mol Cell Proteomics 8: 612–623

Kim HB, Schaller H, Goh CH, Kwon M, Choe S, An CS, Durst F, Feldmann KA, Feyereisen R (2005) *Arabidopsis* cyp51 mutant shows postembryonic seedling lethality associated with lack of membrane integrity. Plant Physiol 138: 2033–2047

Klahre U, Noguchi T, Fujioka S, Takatsuto S, Yokota T, Nomura T, Yoshida S, Chua NH (1998) The *Arabidopsis* DIMINUTO/DWARF1 gene encodes a protein involved in steroid synthesis. Plant Cell 10: 1677–1690

Kribii R, Arró M, Del Arco A, González V, Balcells L, Delourme D, Ferrer A, Karst F, Boronat A (1997) Cloning and characterization of the *Arabidopsis thaliana* SQS1 gene encoding squalene synthase - involvement of the C-terminal region of the enzyme in the channeling of squalene through the sterol pathway. Eur J Biochem 249: 61–69

Kushiro M, Nakano T, Sato K, Yamagishi K, Asami T, Nakano A, Takatsuto S, Fujioka S, Ebizuka Y, Yoshida S (2001) Obtusifoliol 14alpha-demethylase (CYP51) antisense *Arabidopsis* shows slow growth and long life. Biochem Biophys Res Commun 285: 98–104

Lange BM, Rios-Estepa R (2014) Kinetic modeling of plant metabolism and its predictive power: peppermint essential oil biosynthesis as an example. Methods Mol Biol 1083: 287–311

Lange I, Poirier BC, Herron BK, Lange BM (2015) Comprehensive assessment of transcriptional regulation facilitates metabolic engineering of isoprenoid accumulation in Arabidopsis. Plant Physiol 169: 1595–1606

Lecain E, Chenivesse X, Spagnoli R, Pompon D (1996) Cloning by metabolic interference in yeast and enzymatic characterization of *Arabidopsis thaliana* sterol delta 7-reductase. J Biol Chem 271: 10866–10873

Li F, Asami T, Wu X, Tsang EW, Cutler AJ (2007) A putative hydroxysteroid dehydrogenase involved in regulating plant growth and development. Plant Physiol 145: 87–97

Lovato MA, Hart EA, Segura MJ, Giner JL, Matsuda SP (2000) Functional cloning of an *Arabidopsis thaliana* cDNA encoding cycloeucalenol cycloisomerase. J Biol Chem 275: 13394–13397

Men S, Boutté Y, Ikeda Y, Li X, Palme K, Stierhof YD, Hartmann MA, Moritz T, Grebe M (2008) Sterol-dependent endocytosis mediates post cytokinetic acquisition of PIN2 auxin efflux carrier polarity. Nat Cell Biol 10: 237–244

Meng L and Feldman L (2010) A rapid TRIzol-based two-step method for DNA-free RNA extraction from Arabidopsis siliques and dry seeds. Biotechnol J 5: 183–186

Minami A, Tominaga Y, Furuto A, Kondo M, Kawamura Y, Uemura M (2015) Arabidopsis dynamin-related protein 1E in sphingolipid-enriched plasma membrane domains is associated with the development of freezing tolerance. Plant J 83: 501–514

Moreau RA, Nyström L, Whitaker BD, Winkler-Moser JK, Baer DJ, Gebauer SK, Hicks KB (2018) Phytosterols and their derivatives: structural diversity, distribution, metabolism, analysis, and health-promoting uses. Prog Lipid Res. 70: 35–61

Morikawa T, Mizutani M, Ohta D (2006) Cytochrome P450 subfamily CYP710A genes encode sterol C-22 desaturase in plants. Biochem Soc Trans 34: 1202–1205

Nes WD, Song Z, Dennis AL, Zhou W, Nam J, Miller MB (2003) Biosynthesis of phytosterols. Kinetic mechanism for the enzymatic C-methylation of sterols. J Biol Chem 278: 34505–34516

Pascal S, Taton M, Rahier A (1993) Plant sterol biosynthesis. Identification and characterization of two distinct microsomal oxidative enzymatic systems involved in sterol C4-demethylation. J Biol Chem 268: 11639–11654

Patterson GW, Hugly S, Harrison D (1993) Sterols and phytyl esters of Arabidopsis thaliana under normal and chilling temperatures. Phytochemistry 33: 1381–1383

Qian P, Han B, Forestier E, Hu Z, Gao N, Lu W, Schaller H, Li J, Hou S (2013) Sterols are required for cell-fate commitment and maintenance of the stomatal lineage in Arabidopsis. Plant J 74: 1029–1044

Rahier A, Taton M, Benveniste P (1989) Cycloeucalenol-obtusifoliol isomerase. Structural requirements for transformation and binding of substrates and inhibitors. Eur J Biochem 181: 615–626

Rahier A, Darnet S, Bouvier F, Camara B, Bard M (2006) Molecular and enzymatic characterizations of novel bifunctional 3beta-hydroxysteroid dehydrogenases/C-4 decarboxylases from *Arabidopsis thaliana*. J Biol Chem 281: 27264–2777

Rahier A (2011) Dissecting the sterol C-4 demethylation process in higher plants. From structures and genes to catalytic mechanism. Steroids 76: 340–352

Rasbery JM, Shan H, LeClair RJ, Norman M, Matsuda SP, Bartel B (2007) *Arabidopsis thaliana* squalene epoxidase 1 is essential for root and seed development. J Biol Chem 282: 17002–17013

Rios-Estepa R, Turner GW, Lee JM, Croteau RB, Lange BM (2008) A systems biology approach identifies biochemical mechanisms regulating monoterpenoid essential oil composition in peppermint. Proc Natl Acad Sci USA 105: 2818–2823

Rios-Estepa R, Lange I, Lee JM, Lange BM (2010) Mathematical modeling-guided evaluation of biochemical, developmental, environmental, and genotypic determinants of essential oil composition and yield in peppermint leaves. Plant Physiol 152: 2105–2119

Rondet S, Taton M, Rahier A (1999) Identification, characterization, and partial purification of 4 alpha-carboxysterol-C3-dehydrogenase/ C4-decarboxylase from *Zea mays*. Arch Biochem Biophys 366: 249–260

Schmid M, Davison TS, Henz SR, Pape UJ, Demar M, Vingron M, Schölkopf B, Weigel D, Lohmann JU (2005) A gene expression map of Arabidopsis thaliana development. Nat Genet 37: 501–506

Schrick K, Mayer U, Horrichs A, Kuhnt C, Bellini C, Dangl J, Schmidt J, Jürgens G (2000) FACKEL is a sterol C-14 reductase required for organized cell division and expansion in *Arabidopsis* embryogenesis. Genes Dev 14: 1471–1484

Shan H, Wilson WK, Phillips DR, Bartel B, Matsuda SPT (2008) Trinorlupeol, a major nonsterol triterpenoid in Arabidopsis. Organic Lett. 10: 1897–1900

Shi J, Gonzales RA, Bhattacharyya MK (1996) Identification and characterization of an S-adenosyl-L-methionine: delta 24-sterol-C-methyltransferase cDNA from soybean. J Biol Chem 271: 9384–9389

Souter M, Topping J, Pullen M, Friml J, Palme K, Hackett R, Grierson D, Lindsey K (2002) hydra mutants of Arabidopsis are defective in sterol profiles and auxin and ethylene signaling. Plant Cell 14: 1017–1031

Takahashi D, Imai H, Kawamura Y, Uemura M (2016) Lipid profiles of detergent resistant fractions of the plasma membrane in oat and rye in association with cold acclimation and freezing tolerance. Cryobiol 72: 123–134

Taton M, Benveniste P, Rahier A (1989) Microsomal delta 8,14-sterol delta 14-reductase in higher plants. Characterization and inhibition by analogues of a presumptive carbocationic intermediate of the reduction reaction. Eur J Biochem 185: 605–614

Taton M, Rahier A (1991a) Properties and structural requirements for substrate specificity of cytochrome P-450-dependent obtusifoliol 14 alpha-demethylase from maize (*Zea mays*) seedlings. Biochem J 277: 483–492

Taton M, Rahier A (1991b) Identification of delta 5,7-sterol-delta 7-reductase in higher plant microsomes. Biochem Biophys Res Commun 181: 465–473

Taton M, Rahier A. (1996) Plant sterol biosynthesis: identification and characterization of higher plant delta 7-sterol C5(6)-desaturase. Arch Biochem Biophys 325: 279–288

Uemura M, Joseph RA, and Steponkus PL (1995) Cold acclimation of Arabidopsis thaliana; effect on plasma membrane lipid composition and freeze-induced lesions. Plant Physiol 109:15–30

Vaughan MM, Dorothea Tholl D, Tokuhisa JG (2011) An aeroponic culture system for the study of root herbivory on Arabidopsis thaliana. Plant Methods 7: 5

Wewer V, Dombrink I, vom Dorp K, Dörmann P (2011) Quantification of sterol lipids in plants by quadrupole time-of-flight mass spectrometry. J Lipid Res 52: 1039–1054

Willemsen V, Friml J, Grebe M, van den Toorn A, Palme K, Scheres B (2003) Cell polarity and PIN protein positioning in Arabidopsis require *STEROL METHYLTRANSFERASE1* function. Plant Cell 15: 612–625

Zhou W, Nes WD (2003) Sterol methyltransferase2: purification, properties, and inhibition. Arch Biochem Biophys 420: 18–34

