## Supplemental Methods and Data S1 for "Organ-specificity of sterol and triterpene accumulation in *Arabidopsis thaliana*"

**Supplemental Methods and Data File S1**. Organ-specific models of sterol biosynthesis in Arabidopsis (Matlab code).

**Root model – wild-type**

```
function xdot = Root_WT(t,x,E)%it has a flag between x and E

% This function calculates sterol accumulation in Arabidopsis thaliana
% Organ: Roots; Genotype: wild-type
% Michaelis-Menten kinetics are assumed to operate, with the following
% exceptions:

%HMGS is Competitively Inhibited by HMGCoA.
%HMGR is Competitively Inhibited by MVA.
%MK is Competitively Inhibited by FPP.
%SMT1 is Competitively Inhibited by SIT and MeCyA.
%SMT2 is Competitively Inhibited by SIT.

% Kinetic Parameters
%(for details see Table 2 of main manuscript)

% kc units: [1/s]
% KM units: [uM]

kc1 = 2.1    ;
KM1 = 770    ;
kc2 = 0.415  ;
KM2 = 43     ;
kc3= 0.02    ;
KM3= 8.3     ;
kc4= 0.02    ;
KM4= 76      ;
kc5= 0.02    ;
KM5= 42      ;
kc6= 0.02    ;
KM6= 10      ;
kc7= 0.01    ;
KM7= 5.1     ;
kc_7=0.89    ;
KM_7=17      ;
kcipp= 4.397 ;
kcdmapp=1    ;
KMipp = 15.6 ;
KMdmapp = 9  ;
kc9= 0.53    ;
KM9=  9.5    ;
kc10= 0.0183 ;
KM10= 7.7    ;
kc11= 0.02   ;
KM11= 125    ;
kc12= 0.002  ;
KM12= 30     ;
kc13= 0.08   ;   %  SMO1
KM13= 500    ;
kc14= 0.05   ;   %  CECI
KM14= 100    ;
kc15= 0.05   ;   %  CYP51
KM15= 160    ;
kc16= 0.05   ;   %  FACKEL
KM16= 100    ;
kc17= 0.05   ;   %  HYDRA
KM17= 100    ;
kc18= 0.005  ;   %  SMT2/3
KM18= 30     ;
kc19= 0.06   ;   %  DWF7 (campesterol branch)
```

```
KM19= 140    ;
kc20= 0.06   ;   %  DWF5 (campesterol branch)
KM20= 460    ;
kc21= 0.018  ;   %  DWF1 (campesterol branch)
KM21= 150    ;
% There is neither kc22,nor KM22. Further catalysis of CAMP is not considered.
% (flux into brassinosteroids is very low)
kc23= 0.08   ;   %  SMO2 (sitosterol branch & campesterol branch)
KM23= 480    ;
kc24= 0.06   ;   %  DWF7 (sitosterol branch)
KM24= 140    ;
kc25= 0.06   ;   %  DWF5 (sitosterol branch)
KM25= 460    ;
kc26= 0.018  ;   %  DWF1 (sitosterol branch)
KM26= 150    ;
kc27= 0.0005 ;   %  CYP710
KM27= 100    ;

kc40= 0.000001 ;   % Conversion of sterol end products into esters and/or
% integration into membranes

kiHMGS = 9  ;        % Competitive inhibition for HMGS by HMGCoA
kiHMGR = 990;        % Competitive inhibition for HMGR by MVA
kiMK   = 0.1;        % Competitive inhibition for MK by FPP
kiSMT1MeCy = 90;     % Competitive inhibition for SMT1 by 24-methylenecycloartanol
kiSMT1Eyloph = 50;   % Competitive inhibition for SMT1 by ethylidenelophenol
kiSMT1sit = 100;     % Competitive inhibition for SMT1 by sitosterol
kiSMT2sit = 300;     % Competitive inhibition for SMT2 by sitosterol

% Enzyme Concentrations [uM]

E1=0.0583;      %AACT
E2=0.0683;      %HMGS
E3=0.3487;      %HMGR
E4=0.0265;      %MK
E5=0.2530;      %PMK
E6=0.0459;      %MPDC
E7=0.1506;      %IPPI
E8=0.1501;      %FPPS
E9=0.0605;      %SQS
E10=0.1396;     %SQE
E11=0.1389;     %CAS
E12=0.17683;    %SMT1 (main pathway)
E13=0.11107;    %SMO1 (main pathway)
E14=0.0209;     %CECI (main pathway)
E15=0.09899;    %CYP51 (main pathway)
E16=0.04549;    %FACKEL (main pathway)
E17=0.1325;     %HYDRA (main pathway)
E18=0.4065;     %SMT2 for ethylidenelophenol biosynthesis
E19=0.05452;    %DWF7 (campesterol branch)
E20=0.1615;     %DWF5 (campesterol branch)
E21=0.27268;    %DWF1 (campesterol branch)
%There is no E22. Further catalysis of CAMP is not considered.
E23=0.06996;    %SMO2 (sitosterol branch & campesterol branch)
E24=0.05452;    %DWF7 (sitosterol branch)
E25=0.1615;     %DWF5 (sitosterol branch)
E26=0.27268;    %DWF1 (sitosterol branch)
E27=0.01174;    %CYP710A (sitosterol branch)

% Species equations

xdot=[(kc1*E1*x(1)/(x(1)+KM1)-kc1*E1*x(1)/(x(1)+KM1));% Variation of AcCoA
      kc1*E1*x(1)/(x(1)+KM1)-(kc2*E2*x(2)/KM2)/(1+x(2)/KM2+x(3)/kiHMGS); %Variation of AcAcCoA
      (kc2*E2*x(2)/KM2)/(1+x(2)/KM2+x(3)/kiHMGS)-(E3*kc3*x(3)/KM3)/(1+x(3)/KM3 + x(4)/kiHMGR);
%Variation of HMGCoA
      (E3*kc3*x(3)/KM3)/(1+ x(3)/KM3+ x(4)/kiHMGR)- (E4*kc4*x(4)/KM4)/(1+ x(4)/KM4 + x(9)/kiMK);
%Variation of MVA
      (E4*kc4*x(4)/KM4)/(1+ x(4)/KM4 + x(9)/kiMK)- kc5*E5*x(5)/(x(5)+KM5); %Variation of MVAP
```

```
      kc5*E5*x(5)/(x(5)+KM5)-kc6*E6*x(6)/(x(6)+KM6); %Variation of MVAPP
```

```
      kc6*E6*x(6)/(x(6)+KM6)+kc_7*E7*x(8)/(x(8)+KM_7)-kc7*E7*x(7)/(x(7)+KM7)-
(kcipp*E8*KMdmapp*x(7)+kcdmapp*E8*KMipp*x(8))/(KMdmapp*x(7)+KMipp*x(8)+KMipp*KMdmapp); %Variation
of IPP
      kc7*E7*x(7)/(x(7)+KM7)-kc_7*E7*x(8)/(x(8)+KM_7)-
(kcipp*E8*KMdmapp*x(7)+kcdmapp*E8*KMipp*x(8))/(KMdmapp*x(7)+KMipp*x(8)+KMipp*KMdmapp); %Variation
of DMAPP
      0.89*(kcipp*E8*KMdmapp*x(7)+kcdmapp*E8*KMipp*x(8))/(KMdmapp*x(7)+KMipp*x(8)+KMipp*KMdmapp)-
kc9*E9*x(9)/(x(9)+KM9); %Variation of FPP - 9 -
      kc9*E9*x(9)/(x(9)+KM9)-kc10*E10*x(10)/(x(10)+KM10); %Variation of Squalene   - 10 -
      0.97*(kc10*E10*x(10)/(x(10)+KM10))-kc11*E11*x(11)/(x(11)+KM11); %Variation of OxSqualene
0.97
      kc11*E11*x(11)/(x(11)+KM11)-(kc12*E12*x(12))/(KM12*(1 + x(27)/kiSMT1sit + x(13)/kiSMT1MeCy)
+x(23)/kiSMT1Eyloph + x(12));  % Variation of Cycloartenol   -12 -
      (kc12*E12*x(12))/(KM12*(1 + x(27)/kiSMT1sit + x(13)/kiSMT1MeCy) +x(23)/kiSMT1Eyloph +
x(12))- kc13*E13*x(13)/(x(13)+KM13); %Variation in 24-MeCycloartenol - 13 -
      kc13*E13*x(13)/(x(13)+KM13)-kc14*E14*x(14)/(x(14)+KM14); %Variation in Cycloeuclenol - 14 -
      kc14*E14*x(14)/(x(14)+KM14)-kc15*E15*x(15)/(x(15)+KM15); %Variation in Obtusifoliol  - 15 -
      kc15*E15*x(15)/(x(15)+KM15)-kc16*E16*x(16)/(x(16)+KM16); %Variation in CYP51 product  - 16
-
      kc16*E16*x(16)/(x(16)+KM16)-kc17*E17*x(17)/(x(17)+KM17); %Variation in 4-MeFecosterol  - 17
-
      kc17*E17*x(17)/(x(17)+KM17)-(kc23*E23*x(18)/(x(18)+KM23))-(kc18*E18*x(18)/KM18)/(1 +
x(18)/KM18 + x(27)/kiSMT2sit); %Variation in 4-MyLophenol
      (kc23*E23*x(18)/(x(18)+KM23))- (kc19*E19*x(19)/(x(19)+KM19)); %Variation in Episterol
     (kc19*E19*x(19)/(x(19)+KM19))-kc20*E20*x(20)/(x(20)+KM20); %Variation in 5-Dehydroepisterol
-  26 -
      kc20*E20*x(20)/(x(20)+KM20)-kc21*E21*x(21)/(x(21)+KM21); %Variation in 24-MyCholesterol
      kc21*E21*x(21)/(x(21)+KM21)- kc40*x(22); %Variation in Campesterol  - 28 -
      (kc18*E18*x(18)/KM18)/(1 + x(18)/KM18 + x(27)/kiSMT2sit)-kc23*E23*x(23)/(x(23)+KM23);
%Variation in 24-EyLophenol - 19 -
      kc23*E23*x(23)/(x(23)+KM23)-(kc24*E24*x(24)/(x(24)+KM24)); %Variation in Avenasterol
     (kc24*E24*x(24)/(x(24)+KM24))-kc25*E25*x(25)/(x(25)+KM25); %Variation in 5-
Dehydroavenasterol   - 21 -
      kc25*E25*x(25)/(x(25)+KM25)-(kc26*E26*x(26)/(x(26)+KM26)); %Variation in Isofucosterol
      (kc26*E26*x(26)/(x(26)+KM26))-kc27*E27*x(27)/(x(27)+KM27) - kc40*x(27); %********Variation
in Sitosterol - 23 -
      kc27*E27*x(27)/(x(27)+KM27)- kc40*x(28)]; %*******Variation in Stigmasterol  - 24 -
```

**Script file – root – wild-type**

```
%This program calculates the sterol accumulation patterns in Arabidopsis thaliana

%Definition of Variables
%[AcCoA]=x(1), [AcAcCoA]=x(2) , [HMGCoA]=x(3), [MVA]=x(4), [MVAP]=x(5), [MVAPP]=x(6)
%[IPP]=x(7) , [DMAPP]=x(8), [FPP]=x(9),  [SQ]=x(10),  [OxSQ]=x(11), [CyA]=x(12)
%[MeCyA]=x(13), {...

%clear
xdot = zeros(28,1); %Metabolite vector

%Time interval
tspan = [0 4320000];  %[S]  (50 days)

%vector of initial conditions
xdot0 = [0.0053;0;0;0;0;0;0;0;0;0;0;0;0;0;0;0;0;0;0;0;0;0;0;0;0;0;0;0];

%[t,x] = ode15s('Root_WT', tspan, xdot0);
[t,x] = ode15s('Root_WT', tspan, xdot0);

AcCoA=x(:,1);
AcAcCoA=x(:,2);
HMGCoA=x(:,3);
MVA=x(:,4);
MVAP=x(:,5);
MVAPP=x(:,6);
IPP=x(:,7);
DMAPP=x(:,8);
FPP=x(:,9);
SQ=x(:,10);
OxSQ=x(:,11);
```

```
CyA=x(:,12);
MeCyA=x(:,13);
Cycloeucalenol=x(:,14);
Obtusifoliol=x(:,15);
UNKNOWN=x(:,16);
MeFecosterol=x(:,17);
MyLophenol=x(:,18);
Episterol=x(:,19);
Dehydroepisterol=x(:,20);
MyChl=x(:,21);
CAM=x(:,22);
EyLophenol=x(:,23);
Avenasterol=x(:,24);
Dehydroavenasterol=x(:,25);
IFUC=x(:,26);
SIT=x(:,27);
STGM=x(:,28);

%PLOTS

%plot(t/86400,SIT,'b',t/86400,IFUC,'k',t/86400,CyA,'r',t/86400,MeCyA,'c')
plot(t/86400,CyA,'g',t/86400,MeCyA,'k')
hold on
plot(t/86400,CAM,'r','LineWidth',4)
hold on
plot(t/86400,IFUC,'c')
hold on
plot(t/86400,SIT,'b','LineWidth',4)
hold on
plot(t/86400,STGM,'m','LineWidth',4)

%plot(t/86400,SIT,'b',t/86400,IFUC,'k',t/86400,CyA,'r',t/86400,MeCyA,'c',t/86400,OxSQ,'g',
t/86400,CAM,'m', t/86400,AcCoA,'y',t/86400,STGM,'r*')
%plot(t/86400,SIT,'b',t/86400,IFUC,'k',t/86400,CyA,'r',t/86400,MeCyA,'c',t/86400,MyLophenol,'g',t
/86400,EyLophenol,'m',t/86400,MVA,'y',t/86400,OxSQ,'m',t/86400,Cycloeucalenol,'g',t/86400,Obtusif
oliol,'k',t/86400,MeFecosterol,'r',t/86400,Avenasterol,'g',t/86400,Dehydroavenasterol,'g',t/86400
,AcCoA,'g',t/86400,IPP,'r',t/86400,SQ,'r',t/86400,HMGCoA,'g',t/86400,DMAPP,'m')

%legend('SIT','IFUC','CyA','MeCyA','MyLophenol','EyLophenol','MVA','OxSQ') %
legend('CyA','MeCyA','CAM','IFUC','SIT','STGM')

title('Sterol accumulation in Arabidopsis thaliana leaves - HMG1 ')
xlabel('Time (d)')
ylabel('Sterol Concentration (uM/g fresh weight)')
```

**Root model – HMG1**

```
function xdot = Root_HMG1(t,x,E)%it has a flag between x and E

% This function calculates sterol accumulation in Arabidopsis thaliana
% Organ: Roots; Genotype: HMG1
% Michaelis-Menten kinetics are assumed to operate, with the following
% exceptions:

%HMGS is Competitively Inhibited by HMGCoA.
%HMGR is Competitively Inhibited by MVA.
%MK is Competitively Inhibited by FPP.
%SMT1 is Competitively Inhibited by SIT and MeCyA.
%SMT2 is Competitively Inhibited by SIT.

% Kinetic Parameters
%(for details see Table 2 of main manuscript)

% kc units: [1/s]
% KM units: [uM]

kc1 = 2.1    ;
KM1 = 770    ;
kc2 = 0.415  ;
KM2 = 43     ;
kc3= 0.02    ;
KM3= 8.3     ;
kc4= 0.02    ;
KM4= 76      ;
kc5= 0.02    ;
KM5= 42      ;
kc6= 0.02    ;
KM6= 10      ;
kc7= 0.01    ;
KM7= 5.1     ;
kc_7=0.89    ;
KM_7=17      ;
kcipp= 4.397 ;
kcdmapp=1    ;
KMipp = 15.6 ;
KMdmapp = 9  ;
kc9= 0.53    ;
KM9=  9.5    ;
kc10= 0.0183 ;
KM10= 7.7    ;
kc11= 0.02   ;
KM11= 125    ;
kc12= 0.002  ;
KM12= 30     ;
kc13= 0.08   ;   %  SMO1
KM13= 500    ;
kc14= 0.05   ;   %  CECI
KM14= 100    ;
kc15= 0.05   ;   %  CYP51
KM15= 160    ;
kc16= 0.05   ;   %  FACKEL
KM16= 100    ;
kc17= 0.05   ;   %  HYDRA
KM17= 100    ;
kc18= 0.005  ;   %  SMT2/3
KM18= 30     ;
kc19= 0.06   ;   %  DWF7 (campesterol branch)
KM19= 140    ;
kc20= 0.06   ;   %  DWF5 (campesterol branch)
KM20= 460    ;
kc21= 0.018  ;   %  DWF1 (campesterol branch)
KM21= 150    ;
% There is neither kc22,nor KM22. Further catalysis of CAMP is not considered.
% (flux into brassinosteroids is very low)
kc23= 0.08   ;   %  SMO2 (sitosterol branch & campesterol branch)
KM23= 480    ;
```

```
kc24= 0.06   ;   %  DWF7 (sitosterol branch)
KM24= 140    ;
kc25= 0.06   ;   %  DWF5 (sitosterol branch)
KM25= 460    ;
kc26= 0.018  ;   %  DWF1 (sitosterol branch)
KM26= 150    ;
kc27= 0.0005 ;   %  CYP710
KM27= 100    ;

kc40= 0.000003 ;   % Conversion of sterol end products into esters and/or
% integration into membranes

kiHMGS = 9  ;        % Competitive inhibition for HMGS by HMGCoA
kiHMGR = 990;        % Competitive inhibition for HMGR by MVA
kiMK   = 0.1;        % Competitive inhibition for MK by FPP
kiSMT1MeCy = 90;     % Competitive inhibition for SMT1 by 24-methylenecycloartanol
kiSMT1Eyloph = 50;   % Competitive inhibition for SMT1 by ethylidenelophenol
kiSMT1sit = 100;     % Competitive inhibition for SMT1 by sitosterol
kiSMT2sit = 300;     % Competitive inhibition for SMT2 by sitosterol

% Enzyme Concentrations [uM]

E1=0.0583;      %AACT
E2=0.0683;      %HMGS
E3=1.7435;      %HMGR
E4=0.0265;      %MK
E5=0.2530;      %PMK
E6=0.0459;      %MPDC
E7=0.1506;      %IPPI
E8=0.1501;      %FPPS
E9=0.0605;      %SQS
E10=0.1396;     %SQE
E11=0.1389;     %CAS
E12=0.17683;    %SMT1 (main pathway)
E13=0.11107;    %SMO1 (main pathway)
E14=0.0209;     %CECI (main pathway)
E15=0.09899;    %CYP51 (main pathway)
E16=0.04549;    %FACKEL (main pathway)
E17=0.1325;     %HYDRA (main pathway)
E18=0.4065;     %SMT2 for ethylidenelophenol biosynthesis
E19=0.05452;    %DWF7 (campesterol branch)
E20=0.1615;     %DWF5 (campesterol branch)
E21=0.27268;    %DWF1 (campesterol branch)
%There is no E22. Further catalysis of CAMP is not considered.
E23=0.06996;    %SMO2 (sitosterol branch & campesterol branch)
E24=0.05452;    %DWF7 (sitosterol branch)
E25=0.1615;     %DWF5 (sitosterol branch)
E26=0.27268;    %DWF1 (sitosterol branch)
E27=0.01174;    %CYP710A (sitosterol branch)

% Species equations

xdot=[(kc1*E1*x(1)/(x(1)+KM1)-kc1*E1*x(1)/(x(1)+KM1));% Variation of AcCoA
      kc1*E1*x(1)/(x(1)+KM1)-(kc2*E2*x(2)/KM2)/(1+x(2)/KM2+x(3)/kiHMGS); %Variation of AcAcCoA
      (kc2*E2*x(2)/KM2)/(1+x(2)/KM2+x(3)/kiHMGS)-(E3*kc3*x(3)/KM3)/(1+x(3)/KM3 + x(4)/kiHMGR);
%Variation of HMGCoA
      (E3*kc3*x(3)/KM3)/(1+ x(3)/KM3+ x(4)/kiHMGR)- (E4*kc4*x(4)/KM4)/(1+ x(4)/KM4 + x(9)/kiMK);
%Variation of MVA
      (E4*kc4*x(4)/KM4)/(1+ x(4)/KM4 + x(9)/kiMK)- kc5*E5*x(5)/(x(5)+KM5); %Variation of MVAP
- 5 -
      kc5*E5*x(5)/(x(5)+KM5)-kc6*E6*x(6)/(x(6)+KM6); %Variation of MVAPP
      kc6*E6*x(6)/(x(6)+KM6)+kc_7*E7*x(8)/(x(8)+KM_7)-kc7*E7*x(7)/(x(7)+KM7)-
(kcipp*E8*KMdmapp*x(7)+kcdmapp*E8*KMipp*x(8))/(KMdmapp*x(7)+KMipp*x(8)+KMipp*KMdmapp); %Variation
of IPP
      kc7*E7*x(7)/(x(7)+KM7)-kc_7*E7*x(8)/(x(8)+KM_7)-
(kcipp*E8*KMdmapp*x(7)+kcdmapp*E8*KMipp*x(8))/(KMdmapp*x(7)+KMipp*x(8)+KMipp*KMdmapp); %Variation
of DMAPP
      0.89*(kcipp*E8*KMdmapp*x(7)+kcdmapp*E8*KMipp*x(8))/(KMdmapp*x(7)+KMipp*x(8)+KMipp*KMdmapp)-
kc9*E9*x(9)/(x(9)+KM9); %Variation of FPP - 9 -
      kc9*E9*x(9)/(x(9)+KM9)-kc10*E10*x(10)/(x(10)+KM10); %Variation of Squalene   - 10 -
```

```
      0.97*(kc10*E10*x(10)/(x(10)+KM10))-kc11*E11*x(11)/(x(11)+KM11); %Variation of OxSqualene
0.97
      kc11*E11*x(11)/(x(11)+KM11)-(kc12*E12*x(12))/(KM12*(1 + x(27)/kiSMT1sit + x(13)/kiSMT1MeCy)
+x(23)/kiSMT1Eyloph + x(12));  % Variation of Cycloartenol   -12 -
      (kc12*E12*x(12))/(KM12*(1 + x(27)/kiSMT1sit + x(13)/kiSMT1MeCy) +x(23)/kiSMT1Eyloph +
x(12))- kc13*E13*x(13)/(x(13)+KM13); %Variation in 24-MeCycloartenol - 13 -
      kc13*E13*x(13)/(x(13)+KM13)-kc14*E14*x(14)/(x(14)+KM14); %Variation in Cycloeuclenol - 14 -
      kc14*E14*x(14)/(x(14)+KM14)-kc15*E15*x(15)/(x(15)+KM15); %Variation in Obtusifoliol  - 15 -
      kc15*E15*x(15)/(x(15)+KM15)-kc16*E16*x(16)/(x(16)+KM16); %Variation in CYP51 product  - 16
-
      kc16*E16*x(16)/(x(16)+KM16)-kc17*E17*x(17)/(x(17)+KM17); %Variation in 4-MeFecosterol  - 17
-
      kc17*E17*x(17)/(x(17)+KM17)-(kc23*E23*x(18)/(x(18)+KM23))-(kc18*E18*x(18)/KM18)/(1 +
x(18)/KM18 + x(27)/kiSMT2sit); %Variation in 4-MyLophenol
      (kc23*E23*x(18)/(x(18)+KM23))- (kc19*E19*x(19)/(x(19)+KM19)); %Variation in Episterol
     (kc19*E19*x(19)/(x(19)+KM19))-kc20*E20*x(20)/(x(20)+KM20); %Variation in 5-Dehydroepisterol
-  26 -
      kc20*E20*x(20)/(x(20)+KM20)-kc21*E21*x(21)/(x(21)+KM21); %Variation in 24-MyCholesterol
      kc21*E21*x(21)/(x(21)+KM21)- kc40*x(22); %Variation in Campesterol  - 28 -
      (kc18*E18*x(18)/KM18)/(1 + x(18)/KM18 + x(27)/kiSMT2sit)-kc23*E23*x(23)/(x(23)+KM23);
%Variation in 24-EyLophenol - 19 -
      kc23*E23*x(23)/(x(23)+KM23)-(kc24*E24*x(24)/(x(24)+KM24)); %Variation in Avenasterol
     (kc24*E24*x(24)/(x(24)+KM24))-kc25*E25*x(25)/(x(25)+KM25); %Variation in 5-
Dehydroavenasterol   - 21 -
      kc25*E25*x(25)/(x(25)+KM25)-(kc26*E26*x(26)/(x(26)+KM26)); %Variation in Isofucosterol
      (kc26*E26*x(26)/(x(26)+KM26))-kc27*E27*x(27)/(x(27)+KM27) - kc40*x(27); %********Variation
in Sitosterol - 23 -
      kc27*E27*x(27)/(x(27)+KM27)- kc40*x(28)]; %*******Variation in Stigmasterol  - 24 -
```

**Script file – root – HMG1**

```
%This program calculates the sterol accumulation patterns in Arabidopsis thaliana

%Definition of Variables
%[AcCoA]=x(1), [AcAcCoA]=x(2) , [HMGCoA]=x(3), [MVA]=x(4), [MVAP]=x(5), [MVAPP]=x(6)
%[IPP]=x(7) , [DMAPP]=x(8), [FPP]=x(9),  [SQ]=x(10),  [OxSQ]=x(11), [CyA]=x(12)
%[MeCyA]=x(13), {...

%clear
xdot = zeros(28,1); %Metabolite vector

%Time interval
tspan = [0 4320000];  %[S]  (50 days)

%vector of initial conditions
xdot0 = [0.05;0;0;0;0;0;0;0;0;0;0;0;0;0;0;0;0;0;0;0;0;0;0;0;0;0;0;0];

%[t,x] = ode15s('Root_HMG1', tspan, xdot0);
[t,x] = ode15s('Root_HMG1', tspan, xdot0);

AcCoA=x(:,1);
AcAcCoA=x(:,2);
HMGCoA=x(:,3);
MVA=x(:,4);
MVAP=x(:,5);
MVAPP=x(:,6);
IPP=x(:,7);
DMAPP=x(:,8);
FPP=x(:,9);
SQ=x(:,10);
OxSQ=x(:,11);
CyA=x(:,12);
MeCyA=x(:,13);
Cycloeucalenol=x(:,14);
Obtusifoliol=x(:,15);
UNKNOWN=x(:,16);
MeFecosterol=x(:,17);
MyLophenol=x(:,18);
Episterol=x(:,19);
Dehydroepisterol=x(:,20);
```

```
MyChl=x(:,21);
CAM=x(:,22);
EyLophenol=x(:,23);
Avenasterol=x(:,24);
Dehydroavenasterol=x(:,25);
IFUC=x(:,26);
SIT=x(:,27);
STGM=x(:,28);

%PLOTS

%plot(t/86400,SIT,'b',t/86400,IFUC,'k',t/86400,CyA,'r',t/86400,MeCyA,'c')
plot(t/86400,CyA,'g',t/86400,MeCyA,'k')
hold on
plot(t/86400,CAM,'r','LineWidth',4)
hold on
plot(t/86400,IFUC,'c')
hold on
plot(t/86400,SIT,'b','LineWidth',4)
hold on
plot(t/86400,STGM,'m','LineWidth',4)

%plot(t/86400,SIT,'b',t/86400,IFUC,'k',t/86400,CyA,'r',t/86400,MeCyA,'c',t/86400,OxSQ,'g',
t/86400,CAM,'m', t/86400,AcCoA,'y',t/86400,STGM,'r*')
%plot(t/86400,SIT,'b',t/86400,IFUC,'k',t/86400,CyA,'r',t/86400,MeCyA,'c',t/86400,MyLophenol,'g',t
/86400,EyLophenol,'m',t/86400,MVA,'y',t/86400,OxSQ,'m',t/86400,Cycloeucalenol,'g',t/86400,Obtusif
oliol,'k',t/86400,MeFecosterol,'r',t/86400,Avenasterol,'g',t/86400,Dehydroavenasterol,'g',t/86400
,AcCoA,'g',t/86400,IPP,'r',t/86400,SQ,'r',t/86400,HMGCoA,'g',t/86400,DMAPP,'m')

%legend('SIT','IFUC','CyA','MeCyA','MyLophenol','EyLophenol','MVA','OxSQ') %
legend('CyA','MeCyA','CAM','IFUC','SIT','STGM')

title('Sterol accumulation in Arabidopsis thaliana leaves - HMG1 ')
xlabel('Time (d)')
ylabel('Sterol Concentration (uM/g fresh weight)')
```

**Leaf model – wild-type**

```
function xdot = Leaf_WT(t,x,E)%it has a flag between x and E

% This function calculates sterol accumulation in Arabidopsis thaliana
% Organ: Stems; Genotype: wild-type
% Michaelis-Menten kinetics are assumed to operate, with the following
% exceptions:

%HMGS is Competitively Inhibited by HMGCoA.
%HMGR is Competitively Inhibited by MVA.
%MK is Competitively Inhibited by FPP.
%SMT1 is Competitively Inhibited by SIT and MeCyA.
%SMT2 is Competitively Inhibited by SIT.

% Kinetic Parameters
%(for details see Table 2 of main manuscript)

% kc units: [1/s]
% KM units: [uM]

kc1 = 2.1    ;
KM1 = 770    ;
kc2 = 0.415  ;
KM2 = 43     ;
kc3= 0.02    ;
KM3= 8.3     ;
kc4= 0.02    ;
KM4= 76      ;
kc5= 0.02    ;
KM5= 42      ;
kc6= 0.02    ;
KM6= 10      ;
kc7= 0.01    ;
KM7= 5.1     ;
kc_7=0.89    ;
KM_7=17      ;
kcipp= 4.397 ;
kcdmapp=1    ;
KMipp = 15.6 ;
KMdmapp = 9  ;
kc9= 0.53    ;
KM9=  9.5    ;
kc10= 0.0183 ;
KM10= 7.7    ;
kc11= 0.02   ;
KM11= 125    ;
kc12= 0.002  ;
KM12= 30     ;
kc13= 0.08   ;   %  SMO1
KM13= 500    ;
kc14= 0.05   ;   %  CECI
KM14= 100    ;
kc15= 0.05   ;   %  CYP51
KM15= 160    ;
kc16= 0.05   ;   %  FACKEL
KM16= 100    ;
kc17= 0.05   ;   %  HYDRA
KM17= 100    ;
kc18= 0.005  ;   %  SMT2/3
KM18= 30     ;
kc19= 0.06   ;   %  DWF7 (campesterol branch)
KM19= 140    ;
kc20= 0.06   ;   %  DWF5 (campesterol branch)
KM20= 460    ;
kc21= 0.018  ;   %  DWF1 (campesterol branch)
KM21= 150    ;
% There is neither kc22,nor KM22. Further catalysis of CAMP is not considered.
% (flux into brassinosteroids is very low)
kc23= 0.08   ;   %  SMO2 (sitosterol branch & campesterol branch)
KM23= 480    ;
```

```
kc24= 0.06   ;   %  DWF7 (sitosterol branch)
KM24= 140    ;
kc25= 0.06   ;   %  DWF5 (sitosterol branch)
KM25= 460    ;
kc26= 0.018  ;   %  DWF1 (sitosterol branch)
KM26= 150    ;
kc27= 0.0005 ;   %  CYP710
KM27= 100    ;

kc40= 0.000001 ;   % Conversion of sterol end products into esters and/or
% integration into membranes

kiHMGS = 9  ;       % Competitive inhibition for HMGS by HMGCoA
kiHMGR = 990;       % Competitive inhibition for HMGR by MVA
kiMK   = 0.1;       % Competitive inhibition for MK by FPP
kiSMT1MeCy = 90;    % Competitive inhibition for SMT1 by 24-methylenecycloartanol
kiSMT1Eyloph = 50;  % Competitive inhibition for SMT1 by ethylidenelophenol
kiSMT1sit = 100;    % Competitive inhibition for SMT1 by sitosterol
kiSMT2sit = 300;    % Competitive inhibition for SMT2 by sitosterol

% Enzyme Concentrations [uM]

E1=0.0583;      %AACT
E2=0.0683;      %HMGS
E3=0.2352;      %HMGR
E4=0.0265;      %MK
E5=0.2530;      %PMK
E6=0.0459;      %MPDC
E7=0.1506;      %IPPI
E8=0.1501;      %FPPS
E9=0.0605;      %SQS
E10=0.1396;     %SQE
E11=0.1389;     %CAS
E12=0.17683;    %SMT1 (main pathway)
E13=0.11107;    %SMO1 (main pathway)
E14=0.0209;     %CECI (main pathway)
E15=0.09899;    %CYP51 (main pathway)
E16=0.04549;    %FACKEL (main pathway)
E17=0.1325;     %HYDRA (main pathway)
E18=0.2679;     %SMT2 For Ethyllophenol biosynthesis
E19=0.05452;    %DWF7 (campesterol branch)
E20=0.1615;     %DWF5 (campesterol branch)
E21=0.27268;    %DWF1 (campesterol branch)
%There is no E22. Further catalysis of CAMP is not considered.
E23=0.06996;    %SMO2 (sitosterol branch & campesterol branch)
E24=0.05452;    %DWF7 (sitosterol branch)
E25=0.1615;     %DWF5 (sitosterol branch)
E26=0.27268;    %DWF1 (sitosterol branch)
E27=0.01174;    %CYP710A (sitosterol branch)

% Species equations

xdot=[(kc1*E1*x(1)/(x(1)+KM1)-kc1*E1*x(1)/(x(1)+KM1));% Variation of AcCoA
      kc1*E1*x(1)/(x(1)+KM1)-(kc2*E2*x(2)/KM2)/(1+x(2)/KM2+x(3)/kiHMGS); %Variation of AcAcCoA
      (kc2*E2*x(2)/KM2)/(1+x(2)/KM2+x(3)/kiHMGS)-(E3*kc3*x(3)/KM3)/(1+x(3)/KM3 + x(4)/kiHMGR);
%Variation of HMGCoA
      (E3*kc3*x(3)/KM3)/(1+ x(3)/KM3+ x(4)/kiHMGR)- (E4*kc4*x(4)/KM4)/(1+ x(4)/KM4 + x(9)/kiMK);
%Variation of MVA
      (E4*kc4*x(4)/KM4)/(1+ x(4)/KM4 + x(9)/kiMK)- kc5*E5*x(5)/(x(5)+KM5); %Variation of MVAP
- 5 -
      kc5*E5*x(5)/(x(5)+KM5)-kc6*E6*x(6)/(x(6)+KM6); %Variation of MVAPP
      kc6*E6*x(6)/(x(6)+KM6)+kc_7*E7*x(8)/(x(8)+KM_7)-kc7*E7*x(7)/(x(7)+KM7)-
(kcipp*E8*KMdmapp*x(7)+kcdmapp*E8*KMipp*x(8))/(KMdmapp*x(7)+KMipp*x(8)+KMipp*KMdmapp); %Variation
of IPP
      kc7*E7*x(7)/(x(7)+KM7)-kc_7*E7*x(8)/(x(8)+KM_7)-
(kcipp*E8*KMdmapp*x(7)+kcdmapp*E8*KMipp*x(8))/(KMdmapp*x(7)+KMipp*x(8)+KMipp*KMdmapp); %Variation
of DMAPP
      0.89*(kcipp*E8*KMdmapp*x(7)+kcdmapp*E8*KMipp*x(8))/(KMdmapp*x(7)+KMipp*x(8)+KMipp*KMdmapp)-
kc9*E9*x(9)/(x(9)+KM9); %Variation of FPP - 9 -
      kc9*E9*x(9)/(x(9)+KM9)-kc10*E10*x(10)/(x(10)+KM10); %Variation of Squalene   - 10 -
```

```
      0.97*(kc10*E10*x(10)/(x(10)+KM10))-kc11*E11*x(11)/(x(11)+KM11); %Variation of OxSqualene
0.97
      kc11*E11*x(11)/(x(11)+KM11)-(kc12*E12*x(12))/(KM12*(1 + x(27)/kiSMT1sit + x(13)/kiSMT1MeCy)
+x(23)/kiSMT1Eyloph + x(12));  % Variation of Cycloartenol   -12 -
      (kc12*E12*x(12))/(KM12*(1 + x(27)/kiSMT1sit + x(13)/kiSMT1MeCy) +x(23)/kiSMT1Eyloph +
x(12))- kc13*E13*x(13)/(x(13)+KM13); %Variation in 24-MeCycloartenol - 13 -
      kc13*E13*x(13)/(x(13)+KM13)-kc14*E14*x(14)/(x(14)+KM14); %Variation in Cycloeuclenol - 14 -
      kc14*E14*x(14)/(x(14)+KM14)-kc15*E15*x(15)/(x(15)+KM15); %Variation in Obtusifoliol  - 15 -
      kc15*E15*x(15)/(x(15)+KM15)-kc16*E16*x(16)/(x(16)+KM16); %Variation in CYP51 product  - 16
-
      kc16*E16*x(16)/(x(16)+KM16)-kc17*E17*x(17)/(x(17)+KM17); %Variation in 4-MeFecosterol  - 17
-
      kc17*E17*x(17)/(x(17)+KM17)-(kc23*E23*x(18)/(x(18)+KM23))-(kc18*E18*x(18)/KM18)/(1 +
x(18)/KM18 + x(27)/kiSMT2sit); %Variation in 4-MyLophenol
      (kc23*E23*x(18)/(x(18)+KM23))- (kc19*E19*x(19)/(x(19)+KM19)); %Variation in Episterol
     (kc19*E19*x(19)/(x(19)+KM19))-kc20*E20*x(20)/(x(20)+KM20); %Variation in 5-Dehydroepisterol
-  26 -
      kc20*E20*x(20)/(x(20)+KM20)-kc21*E21*x(21)/(x(21)+KM21); %Variation in 24-MyCholesterol
      kc21*E21*x(21)/(x(21)+KM21)- kc40*x(22); %Variation in Campesterol  - 28 -
      (kc18*E18*x(18)/KM18)/(1 + x(18)/KM18 + x(27)/kiSMT2sit)-kc23*E23*x(23)/(x(23)+KM23);
%Variation in 24-EyLophenol - 19 -
      kc23*E23*x(23)/(x(23)+KM23)-(kc24*E24*x(24)/(x(24)+KM24)); %Variation in Avenasterol
     (kc24*E24*x(24)/(x(24)+KM24))-kc25*E25*x(25)/(x(25)+KM25); %Variation in 5-
Dehydroavenasterol   - 21 -
      kc25*E25*x(25)/(x(25)+KM25)-(kc26*E26*x(26)/(x(26)+KM26)); %Variation in Isofucosterol
      (kc26*E26*x(26)/(x(26)+KM26))-kc27*E27*x(27)/(x(27)+KM27) - kc40*x(27); %********Variation
in Sitosterol - 23 -
      kc27*E27*x(27)/(x(27)+KM27)- kc40*x(28)]; %*******Variation in Stigmasterol  - 24 -
```

**Script file – leaf- wild-type**

```
%This program calculates the sterol accumulation patterns in Arabidopsis thaliana

%Definition of Variables
%[AcCoA]=x(1), [AcAcCoA]=x(2) , [HMGCoA]=x(3), [MVA]=x(4), [MVAP]=x(5), [MVAPP]=x(6)
%[IPP]=x(7) , [DMAPP]=x(8), [FPP]=x(9),  [SQ]=x(10),  [OxSQ]=x(11), [CyA]=x(12)
%[MeCyA]=x(13), {...

%clear
xdot = zeros(28,1); %Metabolite vector

%Time interval
tspan = [0 4320000];  %[S]  (50 days)

%vector of initial conditions
xdot0 = [0.006;0;0;0;0;0;0;0;0;0;0;0;0;0;0;0;0;0;0;0;0;0;0;0;0;0;0;0];

%[t,x] = ode15s('Stem_WT', tspan, xdot0);
[t,x] = ode15s('Stem_WT', tspan, xdot0);

AcCoA=x(:,1);
AcAcCoA=x(:,2);
HMGCoA=x(:,3);
MVA=x(:,4);
MVAP=x(:,5);
MVAPP=x(:,6);
IPP=x(:,7);
DMAPP=x(:,8);
FPP=x(:,9);
SQ=x(:,10);
OxSQ=x(:,11);
CyA=x(:,12);
MeCyA=x(:,13);
Cycloeucalenol=x(:,14);
Obtusifoliol=x(:,15);
UNKNOWN=x(:,16);
MeFecosterol=x(:,17);
MyLophenol=x(:,18);
Episterol=x(:,19);
Dehydroepisterol=x(:,20);
```

```
MyChl=x(:,21);
CAM=x(:,22);
EyLophenol=x(:,23);
Avenasterol=x(:,24);
Dehydroavenasterol=x(:,25);
IFUC=x(:,26);
SIT=x(:,27);
STGM=x(:,28);

%PLOTS

%plot(t/86400,SIT,'b',t/86400,IFUC,'k',t/86400,CyA,'r',t/86400,MeCyA,'c')
plot(t/86400,CyA,'g',t/86400,MeCyA,'k')
hold on
plot(t/86400,CAM,'r','LineWidth',4)
hold on
plot(t/86400,IFUC,'c')
hold on
plot(t/86400,SIT,'b','LineWidth',4)
hold on
plot(t/86400,STGM,'m','LineWidth',4)

%plot(t/86400,SIT,'b',t/86400,IFUC,'k',t/86400,CyA,'r',t/86400,MeCyA,'c',t/86400,OxSQ,'g',
t/86400,CAM,'m', t/86400,AcCoA,'y',t/86400,STGM,'r*')
%plot(t/86400,SIT,'b',t/86400,IFUC,'k',t/86400,CyA,'r',t/86400,MeCyA,'c',t/86400,MyLophenol,'g',t
/86400,EyLophenol,'m',t/86400,MVA,'y',t/86400,OxSQ,'m',t/86400,Cycloeucalenol,'g',t/86400,Obtusif
oliol,'k',t/86400,MeFecosterol,'r',t/86400,Avenasterol,'g',t/86400,Dehydroavenasterol,'g',t/86400
,AcCoA,'g',t/86400,IPP,'r',t/86400,SQ,'r',t/86400,HMGCoA,'g',t/86400,DMAPP,'m')

%legend('SIT','IFUC','CyA','MeCyA','MyLophenol','EyLophenol','MVA','OxSQ') %
legend('CyA','MeCyA','CAM','IFUC','SIT','STGM')

title('Sterol accumulation in Arabidopsis thaliana leaves - HMG1 ')
xlabel('Time (d)')
ylabel('Sterol Concentration (uM/g fresh weight)')
```

**Leaf model – HMG1**

```
function xdot = Leaf_HMG1(t,x,E)%it has a flag between x and E

% This function calculates sterol accumulation in Arabidopsis thaliana
% Organ: Leaves; Genotype: HMG1
% Michaelis-Menten kinetics are assumed to operate, with the following
% exceptions:

%HMGS is Competitively Inhibited by HMGCoA.
%HMGR is Competitively Inhibited by MVA.
%MK is Competitively Inhibited by FPP.
%SMT1 is Competitively Inhibited by SIT and MeCyA.
%SMT2 is Competitively Inhibited by SIT.

% Kinetic Parameters
%(for details see Table 2 of main manuscript)

% kc units: [1/s]
% KM units: [uM]

kc1 = 2.1    ;
KM1 = 770    ;
kc2 = 0.415  ;
KM2 = 43     ;
kc3= 0.02    ;
KM3= 8.3     ;
kc4= 0.02    ;
KM4= 76      ;
kc5= 0.02    ;
KM5= 42      ;
kc6= 0.02    ;
KM6= 10      ;
kc7= 0.01    ;
KM7= 5.1     ;
kc_7=0.89    ;
KM_7=17      ;
kcipp= 4.397 ;
kcdmapp=1    ;
KMipp = 15.6 ;
KMdmapp = 9  ;
kc9= 0.53    ;
KM9=  9.5    ;
kc10= 0.0183 ;
KM10= 7.7    ;
kc11= 0.02   ;
KM11= 125    ;
kc12= 0.002  ;
KM12= 30     ;
kc13= 0.08   ;   %  SMO1
KM13= 500    ;
kc14= 0.05   ;   %  CECI
KM14= 100    ;
kc15= 0.05   ;   %  CYP51
KM15= 160    ;
kc16= 0.05   ;   %  FACKEL
KM16= 100    ;
kc17= 0.05   ;   %  HYDRA
KM17= 100    ;
kc18= 0.005  ;   %  SMT2/3
KM18= 30     ;
kc19= 0.06   ;   %  DWF7 (campesterol branch)
KM19= 140    ;
kc20= 0.06   ;   %  DWF5 (campesterol branch)
KM20= 460    ;
kc21= 0.018  ;   %  DWF1 (campesterol branch)
KM21= 150    ;
% There is neither kc22,nor KM22. Further catalysis of CAMP is not considered.
% (flux into brassinosteroids is very low)
kc23= 0.08   ;   %  SMO2 (sitosterol branch & campesterol branch)
KM23= 480    ;
```

```
kc24= 0.06   ;   %  DWF7 (sitosterol branch)
KM24= 140    ;
kc25= 0.06   ;   %  DWF5 (sitosterol branch)
KM25= 460    ;
kc26= 0.018  ;   %  DWF1 (sitosterol branch)
KM26= 150    ;
kc27= 0.0005 ;   %  CYP710
KM27= 100    ;

kc40= 0.000002 ;   % Conversion of sterol end products into esters and/or
% integration into membranes

kiHMGS = 9  ;       % Competitive inhibition for HMGS by HMGCoA
kiHMGR = 990;       % Competitive inhibition for HMGR by MVA
kiMK   = 0.1;       % Competitive inhibition for MK by FPP
kiSMT1MeCy = 90;    % Competitive inhibition for SMT1 by 24-methylenecycloartanol
kiSMT1Eyloph = 50;  % Competitive inhibition for SMT1 by ethylidenelophenol
kiSMT1sit = 100;    % Competitive inhibition for SMT1 by sitosterol
kiSMT2sit = 300;    % Competitive inhibition for SMT2 by sitosterol

% Enzyme Concentrations [uM]

E1=0.0583;      %AACT
E2=0.0683;      %HMGS
E3=1.2230;      %HMGR
E4=0.0265;      %MK
E5=0.2530;      %PMK
E6=0.0459;      %MPDC
E7=0.1506;      %IPPI
E8=0.1501;      %FPPS
E9=0.0605;      %SQS
E10=0.1396;     %SQE
E11=0.1389;     %CAS
E12=0.17683;    %SMT1 (main pathway)
E13=0.11107;    %SMO1 (main pathway)
E14=0.0209;     %CECI (main pathway)
E15=0.09899;    %CYP51 (main pathway)
E16=0.04549;    %FACKEL (main pathway)
E17=0.1325;     %HYDRA (main pathway)
E18=0.2679;     %SMT2 For Ethyllophenol biosynthesis
E19=0.05452;    %DWF7 (campesterol branch)
E20=0.1615;     %DWF5 (campesterol branch)
E21=0.27268;    %DWF1 (campesterol branch)
%There is no E22. Further catalysis of CAMP is not considered.
E23=0.06996;    %SMO2 (sitosterol branch & campesterol branch)
E24=0.05452;    %DWF7 (sitosterol branch)
E25=0.1615;     %DWF5 (sitosterol branch)
E26=0.27268;    %DWF1 (sitosterol branch)
E27=0.01174;    %CYP710A (sitosterol branch)

% Species equations

xdot=[(kc1*E1*x(1)/(x(1)+KM1)-kc1*E1*x(1)/(x(1)+KM1));% Variation of AcCoA
      kc1*E1*x(1)/(x(1)+KM1)-(kc2*E2*x(2)/KM2)/(1+x(2)/KM2+x(3)/kiHMGS); %Variation of AcAcCoA
      (kc2*E2*x(2)/KM2)/(1+x(2)/KM2+x(3)/kiHMGS)-(E3*kc3*x(3)/KM3)/(1+x(3)/KM3 + x(4)/kiHMGR);
%Variation of HMGCoA
      (E3*kc3*x(3)/KM3)/(1+ x(3)/KM3+ x(4)/kiHMGR)- (E4*kc4*x(4)/KM4)/(1+ x(4)/KM4 + x(9)/kiMK);
%Variation of MVA
      (E4*kc4*x(4)/KM4)/(1+ x(4)/KM4 + x(9)/kiMK)- kc5*E5*x(5)/(x(5)+KM5); %Variation of MVAP
- 5 -
      kc5*E5*x(5)/(x(5)+KM5)-kc6*E6*x(6)/(x(6)+KM6); %Variation of MVAPP
      kc6*E6*x(6)/(x(6)+KM6)+kc_7*E7*x(8)/(x(8)+KM_7)-kc7*E7*x(7)/(x(7)+KM7)-
(kcipp*E8*KMdmapp*x(7)+kcdmapp*E8*KMipp*x(8))/(KMdmapp*x(7)+KMipp*x(8)+KMipp*KMdmapp); %Variation
of IPP
      kc7*E7*x(7)/(x(7)+KM7)-kc_7*E7*x(8)/(x(8)+KM_7)-
(kcipp*E8*KMdmapp*x(7)+kcdmapp*E8*KMipp*x(8))/(KMdmapp*x(7)+KMipp*x(8)+KMipp*KMdmapp); %Variation
of DMAPP
      0.89*(kcipp*E8*KMdmapp*x(7)+kcdmapp*E8*KMipp*x(8))/(KMdmapp*x(7)+KMipp*x(8)+KMipp*KMdmapp)-
kc9*E9*x(9)/(x(9)+KM9); %Variation of FPP - 9 -
      kc9*E9*x(9)/(x(9)+KM9)-kc10*E10*x(10)/(x(10)+KM10); %Variation of Squalene   - 10 -
```

```
      0.97*(kc10*E10*x(10)/(x(10)+KM10))-kc11*E11*x(11)/(x(11)+KM11); %Variation of OxSqualene
0.97
      kc11*E11*x(11)/(x(11)+KM11)-(kc12*E12*x(12))/(KM12*(1 + x(27)/kiSMT1sit + x(13)/kiSMT1MeCy)
+x(23)/kiSMT1Eyloph + x(12));  % Variation of Cycloartenol   -12 -
      (kc12*E12*x(12))/(KM12*(1 + x(27)/kiSMT1sit + x(13)/kiSMT1MeCy) +x(23)/kiSMT1Eyloph +
x(12))- kc13*E13*x(13)/(x(13)+KM13); %Variation in 24-MeCycloartenol - 13 -
      kc13*E13*x(13)/(x(13)+KM13)-kc14*E14*x(14)/(x(14)+KM14); %Variation in Cycloeuclenol - 14 -
      kc14*E14*x(14)/(x(14)+KM14)-kc15*E15*x(15)/(x(15)+KM15); %Variation in Obtusifoliol  - 15 -
      kc15*E15*x(15)/(x(15)+KM15)-kc16*E16*x(16)/(x(16)+KM16); %Variation in CYP51 product  - 16
-
      kc16*E16*x(16)/(x(16)+KM16)-kc17*E17*x(17)/(x(17)+KM17); %Variation in 4-MeFecosterol  - 17
-
      kc17*E17*x(17)/(x(17)+KM17)-(kc23*E23*x(18)/(x(18)+KM23))-(kc18*E18*x(18)/KM18)/(1 +
x(18)/KM18 + x(27)/kiSMT2sit); %Variation in 4-MyLophenol
      (kc23*E23*x(18)/(x(18)+KM23))- (kc19*E19*x(19)/(x(19)+KM19)); %Variation in Episterol
     (kc19*E19*x(19)/(x(19)+KM19))-kc20*E20*x(20)/(x(20)+KM20); %Variation in 5-Dehydroepisterol
-  26 -
      kc20*E20*x(20)/(x(20)+KM20)-kc21*E21*x(21)/(x(21)+KM21); %Variation in 24-MyCholesterol
      kc21*E21*x(21)/(x(21)+KM21)- kc40*x(22); %Variation in Campesterol  - 28 -
      (kc18*E18*x(18)/KM18)/(1 + x(18)/KM18 + x(27)/kiSMT2sit)-kc23*E23*x(23)/(x(23)+KM23);
%Variation in 24-EyLophenol - 19 -
      kc23*E23*x(23)/(x(23)+KM23)-(kc24*E24*x(24)/(x(24)+KM24)); %Variation in Avenasterol
     (kc24*E24*x(24)/(x(24)+KM24))-kc25*E25*x(25)/(x(25)+KM25); %Variation in 5-
Dehydroavenasterol   - 21 -
      kc25*E25*x(25)/(x(25)+KM25)-(kc26*E26*x(26)/(x(26)+KM26)); %Variation in Isofucosterol
      (kc26*E26*x(26)/(x(26)+KM26))-kc27*E27*x(27)/(x(27)+KM27) - kc40*x(27); %********Variation
in Sitosterol - 23 -
      kc27*E27*x(27)/(x(27)+KM27)- kc40*x(28)]; %*******Variation in Stigmasterol  - 24 -
```

**Script file – leaf – HMG1**

```
%This program calculates the sterol accumulation patterns in Arabidopsis thaliana

%Definition of Variables
%[AcCoA]=x(1), [AcAcCoA]=x(2) , [HMGCoA]=x(3), [MVA]=x(4), [MVAP]=x(5), [MVAPP]=x(6)
%[IPP]=x(7) , [DMAPP]=x(8), [FPP]=x(9),  [SQ]=x(10),  [OxSQ]=x(11), [CyA]=x(12)
%[MeCyA]=x(13), {...

%clear
xdot = zeros(28,1); %Metabolite vector

%Time interval
tspan = [0 4320000];  %[S]  (50 days)

%vector of initial conditions
xdot0 = [0.024;0;0;0;0;0;0;0;0;0;0;0;0;0;0;0;0;0;0;0;0;0;0;0;0;0;0;0];

%[t,x] = ode15s('Leaf_HMG1', tspan, xdot0);
[t,x] = ode15s('Leaf_HMG1', tspan, xdot0);

AcCoA=x(:,1);
AcAcCoA=x(:,2);
HMGCoA=x(:,3);
MVA=x(:,4);
MVAP=x(:,5);
MVAPP=x(:,6);
IPP=x(:,7);
DMAPP=x(:,8);
FPP=x(:,9);
SQ=x(:,10);
OxSQ=x(:,11);
CyA=x(:,12);
MeCyA=x(:,13);
Cycloeucalenol=x(:,14);
Obtusifoliol=x(:,15);
UNKNOWN=x(:,16);
MeFecosterol=x(:,17);
MyLophenol=x(:,18);
Episterol=x(:,19);
Dehydroepisterol=x(:,20);
```

```
MyChl=x(:,21);
CAM=x(:,22);
EyLophenol=x(:,23);
Avenasterol=x(:,24);
Dehydroavenasterol=x(:,25);
IFUC=x(:,26);
SIT=x(:,27);
STGM=x(:,28);

%PLOTS

%plot(t/86400,SIT,'b',t/86400,IFUC,'k',t/86400,CyA,'r',t/86400,MeCyA,'c')
plot(t/86400,CyA,'g',t/86400,MeCyA,'k')
hold on
plot(t/86400,CAM,'r','LineWidth',4)
hold on
plot(t/86400,IFUC,'c')
hold on
plot(t/86400,SIT,'b','LineWidth',4)
hold on
plot(t/86400,STGM,'m','LineWidth',4)

%plot(t/86400,SIT,'b',t/86400,IFUC,'k',t/86400,CyA,'r',t/86400,MeCyA,'c',t/86400,OxSQ,'g',
t/86400,CAM,'m', t/86400,AcCoA,'y',t/86400,STGM,'r*')
%plot(t/86400,SIT,'b',t/86400,IFUC,'k',t/86400,CyA,'r',t/86400,MeCyA,'c',t/86400,MyLophenol,'g',t
/86400,EyLophenol,'m',t/86400,MVA,'y',t/86400,OxSQ,'m',t/86400,Cycloeucalenol,'g',t/86400,Obtusif
oliol,'k',t/86400,MeFecosterol,'r',t/86400,Avenasterol,'g',t/86400,Dehydroavenasterol,'g',t/86400
,AcCoA,'g',t/86400,IPP,'r',t/86400,SQ,'r',t/86400,HMGCoA,'g',t/86400,DMAPP,'m')

%legend('SIT','IFUC','CyA','MeCyA','MyLophenol','EyLophenol','MVA','OxSQ') %
legend('CyA','MeCyA','CAM','IFUC','SIT','STGM')

title('Sterol accumulation in Arabidopsis thaliana leaves - HMG1 ')
xlabel('Time (d)')
ylabel('Sterol Concentration (uM/g fresh weight)')
```

**Stem model – wild-type**

```
function xdot = Stem_WT(t,x,E)%it has a flag between x and E

% This function calculates sterol accumulation in Arabidopsis thaliana
% Organ: Stems; Genotype: wild-type
% Michaelis-Menten kinetics are assumed to operate, with the following
% exceptions:

%HMGS is Competitively Inhibited by HMGCoA.
%HMGR is Competitively Inhibited by MVA.
%MK is Competitively Inhibited by FPP.
%SMT1 is Competitively Inhibited by SIT and MeCyA.
%SMT2 is Competitively Inhibited by SIT.

% Kinetic Parameters
%(for details see Table 2 of main manuscript)

% kc units: [1/s]
% KM units: [uM]

kc1 = 2.1    ;
KM1 = 770    ;
kc2 = 0.415  ;
KM2 = 43     ;
kc3= 0.02    ;
KM3= 8.3     ;
kc4= 0.02    ;
KM4= 76      ;
kc5= 0.02    ;
KM5= 42      ;
kc6= 0.02    ;
KM6= 10      ;
kc7= 0.01    ;
KM7= 5.1     ;
kc_7=0.89    ;
KM_7=17      ;
kcipp= 4.397 ;
kcdmapp=1    ;
KMipp = 15.6 ;
KMdmapp = 9  ;
kc9= 0.53    ;
KM9=  9.5    ;
kc10= 0.0183 ;
KM10= 7.7    ;
kc11= 0.02   ;
KM11= 125    ;
kc12= 0.002  ;
KM12= 30     ;
kc13= 0.08   ;   %  SMO1
KM13= 500    ;
kc14= 0.05   ;   %  CECI
KM14= 100    ;
kc15= 0.05   ;   %  CYP51
KM15= 160    ;
kc16= 0.05   ;   %  FACKEL
KM16= 100    ;
kc17= 0.05   ;   %  HYDRA
KM17= 100    ;
kc18= 0.005  ;   %  SMT2/3
KM18= 30     ;
kc19= 0.06   ;   %  DWF7 (campesterol branch)
KM19= 140    ;
kc20= 0.06   ;   %  DWF5 (campesterol branch)
KM20= 460    ;
kc21= 0.018  ;   %  DWF1 (campesterol branch)
KM21= 150    ;
% There is neither kc22,nor KM22. Further catalysis of CAMP is not considered.
% (flux into brassinosteroids is very low)
kc23= 0.08   ;   %  SMO2 (sitosterol branch & campesterol branch)
KM23= 480    ;
```

```
kc24= 0.06   ;   %  DWF7 (sitosterol branch)
KM24= 140    ;
kc25= 0.06   ;   %  DWF5 (sitosterol branch)
KM25= 460    ;
kc26= 0.018  ;   %  DWF1 (sitosterol branch)
KM26= 150    ;
kc27= 0.0005 ;   %  CYP710
KM27= 100    ;

kc40= 0.000001 ;   % Conversion of sterol end products into esters and/or
% integration into membranes

kiHMGS = 9  ;       % Competitive inhibition for HMGS by HMGCoA
kiHMGR = 990;       % Competitive inhibition for HMGR by MVA
kiMK   = 0.1;       % Competitive inhibition for MK by FPP
kiSMT1MeCy = 90;    % Competitive inhibition for SMT1 by 24-methylenecycloartanol
kiSMT1Eyloph = 50;  % Competitive inhibition for SMT1 by ethylidenelophenol
kiSMT1sit = 100;    % Competitive inhibition for SMT1 by sitosterol
kiSMT2sit = 300;    % Competitive inhibition for SMT2 by sitosterol

% Enzyme Concentrations [uM]

E1=0.0583;      %AACT
E2=0.0683;      %HMGS
E3=0.2393;      %HMGR
E4=0.0265;      %MK
E5=0.2530;      %PMK
E6=0.0459;      %MPDC
E7=0.1506;      %IPPI
E8=0.1501;      %FPPS
E9=0.0605;      %SQS
E10=0.1396;     %SQE
E11=0.1389;     %CAS
E12=0.17683;    %SMT1 (main pathway)
E13=0.11107;    %SMO1 (main pathway)
E14=0.0209;     %CECI (main pathway)
E15=0.09899;    %CYP51 (main pathway)
E16=0.04549;    %FACKEL (main pathway)
E17=0.1325;     %HYDRA (main pathway)
E18=0.2500;     %SMT2 For Ethyllophenol biosynthesis
E19=0.05452;    %DWF7 (campesterol branch)
E20=0.1615;     %DWF5 (campesterol branch)
E21=0.27268;    %DWF1 (campesterol branch)
%There is no E22. Further catalysis of CAMP is not considered.
E23=0.06996;    %SMO2 (sitosterol branch & campesterol branch)
E24=0.05452;    %DWF7 (sitosterol branch)
E25=0.1615;     %DWF5 (sitosterol branch)
E26=0.27268;    %DWF1 (sitosterol branch)
E27=0.01174;    %CYP710A (sitosterol branch)

% Species equations

xdot=[(kc1*E1*x(1)/(x(1)+KM1)-kc1*E1*x(1)/(x(1)+KM1));% Variation of AcCoA
      kc1*E1*x(1)/(x(1)+KM1)-(kc2*E2*x(2)/KM2)/(1+x(2)/KM2+x(3)/kiHMGS); %Variation of AcAcCoA
      (kc2*E2*x(2)/KM2)/(1+x(2)/KM2+x(3)/kiHMGS)-(E3*kc3*x(3)/KM3)/(1+x(3)/KM3 + x(4)/kiHMGR);
%Variation of HMGCoA
      (E3*kc3*x(3)/KM3)/(1+ x(3)/KM3+ x(4)/kiHMGR)- (E4*kc4*x(4)/KM4)/(1+ x(4)/KM4 + x(9)/kiMK);
%Variation of MVA
      (E4*kc4*x(4)/KM4)/(1+ x(4)/KM4 + x(9)/kiMK)- kc5*E5*x(5)/(x(5)+KM5); %Variation of MVAP
- 5 -
      kc5*E5*x(5)/(x(5)+KM5)-kc6*E6*x(6)/(x(6)+KM6); %Variation of MVAPP
      kc6*E6*x(6)/(x(6)+KM6)+kc_7*E7*x(8)/(x(8)+KM_7)-kc7*E7*x(7)/(x(7)+KM7)-
(kcipp*E8*KMdmapp*x(7)+kcdmapp*E8*KMipp*x(8))/(KMdmapp*x(7)+KMipp*x(8)+KMipp*KMdmapp); %Variation
of IPP
      kc7*E7*x(7)/(x(7)+KM7)-kc_7*E7*x(8)/(x(8)+KM_7)-
(kcipp*E8*KMdmapp*x(7)+kcdmapp*E8*KMipp*x(8))/(KMdmapp*x(7)+KMipp*x(8)+KMipp*KMdmapp); %Variation
of DMAPP
      0.89*(kcipp*E8*KMdmapp*x(7)+kcdmapp*E8*KMipp*x(8))/(KMdmapp*x(7)+KMipp*x(8)+KMipp*KMdmapp)-
kc9*E9*x(9)/(x(9)+KM9); %Variation of FPP - 9 -
      kc9*E9*x(9)/(x(9)+KM9)-kc10*E10*x(10)/(x(10)+KM10); %Variation of Squalene   - 10 -
```

```
      0.97*(kc10*E10*x(10)/(x(10)+KM10))-kc11*E11*x(11)/(x(11)+KM11); %Variation of OxSqualene
0.97
      kc11*E11*x(11)/(x(11)+KM11)-(kc12*E12*x(12))/(KM12*(1 + x(27)/kiSMT1sit + x(13)/kiSMT1MeCy)
+x(23)/kiSMT1Eyloph + x(12));  % Variation of Cycloartenol   -12 -
      (kc12*E12*x(12))/(KM12*(1 + x(27)/kiSMT1sit + x(13)/kiSMT1MeCy) +x(23)/kiSMT1Eyloph +
x(12))- kc13*E13*x(13)/(x(13)+KM13); %Variation in 24-MeCycloartenol - 13 -
      kc13*E13*x(13)/(x(13)+KM13)-kc14*E14*x(14)/(x(14)+KM14); %Variation in Cycloeuclenol - 14 -
      kc14*E14*x(14)/(x(14)+KM14)-kc15*E15*x(15)/(x(15)+KM15); %Variation in Obtusifoliol  - 15 -
      kc15*E15*x(15)/(x(15)+KM15)-kc16*E16*x(16)/(x(16)+KM16); %Variation in CYP51 product  - 16
-
      kc16*E16*x(16)/(x(16)+KM16)-kc17*E17*x(17)/(x(17)+KM17); %Variation in 4-MeFecosterol  - 17
-
      kc17*E17*x(17)/(x(17)+KM17)-(kc23*E23*x(18)/(x(18)+KM23))-(kc18*E18*x(18)/KM18)/(1 +
x(18)/KM18 + x(27)/kiSMT2sit); %Variation in 4-MyLophenol
      (kc23*E23*x(18)/(x(18)+KM23))- (kc19*E19*x(19)/(x(19)+KM19)); %Variation in Episterol
     (kc19*E19*x(19)/(x(19)+KM19))-kc20*E20*x(20)/(x(20)+KM20); %Variation in 5-Dehydroepisterol
-  26 -
      kc20*E20*x(20)/(x(20)+KM20)-kc21*E21*x(21)/(x(21)+KM21); %Variation in 24-MyCholesterol
      kc21*E21*x(21)/(x(21)+KM21)- kc40*x(22); %Variation in Campesterol  - 28 -
      (kc18*E18*x(18)/KM18)/(1 + x(18)/KM18 + x(27)/kiSMT2sit)-kc23*E23*x(23)/(x(23)+KM23);
%Variation in 24-EyLophenol - 19 -
      kc23*E23*x(23)/(x(23)+KM23)-(kc24*E24*x(24)/(x(24)+KM24)); %Variation in Avenasterol
     (kc24*E24*x(24)/(x(24)+KM24))-kc25*E25*x(25)/(x(25)+KM25); %Variation in 5-
Dehydroavenasterol   - 21 -
      kc25*E25*x(25)/(x(25)+KM25)-(kc26*E26*x(26)/(x(26)+KM26)); %Variation in Isofucosterol
      (kc26*E26*x(26)/(x(26)+KM26))-kc27*E27*x(27)/(x(27)+KM27) - kc40*x(27); %********Variation
in Sitosterol - 23 -
      kc27*E27*x(27)/(x(27)+KM27)- kc40*x(28)]; %*******Variation in Stigmasterol  - 24 -
```

**Script file – stem – wild-type**

```
%This program calculates the sterol accumulation patterns in Arabidopsis thaliana

%Definition of Variables
%[AcCoA]=x(1), [AcAcCoA]=x(2) , [HMGCoA]=x(3), [MVA]=x(4), [MVAP]=x(5), [MVAPP]=x(6)
%[IPP]=x(7) , [DMAPP]=x(8), [FPP]=x(9),  [SQ]=x(10),  [OxSQ]=x(11), [CyA]=x(12)
%[MeCyA]=x(13), {...

%clear
xdot = zeros(28,1); %Metabolite vector

%Time interval
tspan = [0 4320000];  %[S]  (50 days)

%vector of initial conditions
xdot0 = [0.013;0;0;0;0;0;0;0;0;0;0;0;0;0;0;0;0;0;0;0;0;0;0;0;0;0;0;0];

%[t,x] = ode15s('Stem_WT', tspan, xdot0);
[t,x] = ode15s('Stem_WT', tspan, xdot0);

AcCoA=x(:,1);
AcAcCoA=x(:,2);
HMGCoA=x(:,3);
MVA=x(:,4);
MVAP=x(:,5);
MVAPP=x(:,6);
IPP=x(:,7);
DMAPP=x(:,8);
FPP=x(:,9);
SQ=x(:,10);
OxSQ=x(:,11);
CyA=x(:,12);
MeCyA=x(:,13);
Cycloeucalenol=x(:,14);
Obtusifoliol=x(:,15);
UNKNOWN=x(:,16);
MeFecosterol=x(:,17);
MyLophenol=x(:,18);
Episterol=x(:,19);
Dehydroepisterol=x(:,20);
```

```
MyChl=x(:,21);
CAM=x(:,22);
EyLophenol=x(:,23);
Avenasterol=x(:,24);
Dehydroavenasterol=x(:,25);
IFUC=x(:,26);
SIT=x(:,27);
STGM=x(:,28);

%PLOTS

%plot(t/86400,SIT,'b',t/86400,IFUC,'k',t/86400,CyA,'r',t/86400,MeCyA,'c')
plot(t/86400,CyA,'g',t/86400,MeCyA,'k')
hold on
plot(t/86400,CAM,'r','LineWidth',4)
hold on
plot(t/86400,IFUC,'c')
hold on
plot(t/86400,SIT,'b','LineWidth',4)
hold on
plot(t/86400,STGM,'m','LineWidth',4)

%plot(t/86400,SIT,'b',t/86400,IFUC,'k',t/86400,CyA,'r',t/86400,MeCyA,'c',t/86400,OxSQ,'g',
t/86400,CAM,'m', t/86400,AcCoA,'y',t/86400,STGM,'r*')
%plot(t/86400,SIT,'b',t/86400,IFUC,'k',t/86400,CyA,'r',t/86400,MeCyA,'c',t/86400,MyLophenol,'g',t
/86400,EyLophenol,'m',t/86400,MVA,'y',t/86400,OxSQ,'m',t/86400,Cycloeucalenol,'g',t/86400,Obtusif
oliol,'k',t/86400,MeFecosterol,'r',t/86400,Avenasterol,'g',t/86400,Dehydroavenasterol,'g',t/86400
,AcCoA,'g',t/86400,IPP,'r',t/86400,SQ,'r',t/86400,HMGCoA,'g',t/86400,DMAPP,'m')

%legend('SIT','IFUC','CyA','MeCyA','MyLophenol','EyLophenol','MVA','OxSQ') %
legend('CyA','MeCyA','CAM','IFUC','SIT','STGM')

title('Sterol accumulation in Arabidopsis thaliana leaves - HMG1 ')
xlabel('Time (d)')
ylabel('Sterol Concentration (uM/g fresh weight)')
```

**Stem model – HMG1**

```
function xdot = Stem_HMG1(t,x,E)%it has a flag between x and E

% This function calculates sterol accumulation in Arabidopsis thaliana
% Organ: Leaves; Genotype: HMG1
% Michaelis-Menten kinetics are assumed to operate, with the following
% exceptions:

%HMGS is Competitively Inhibited by HMGCoA.
%HMGR is Competitively Inhibited by MVA.
%MK is Competitively Inhibited by FPP.
%SMT1 is Competitively Inhibited by SIT and MeCyA.
%SMT2 is Competitively Inhibited by SIT.

% Kinetic Parameters
%(for details see Table 2 of main manuscript)

% kc units: [1/s]
% KM units: [uM]

kc1 = 2.1    ;
KM1 = 770    ;
kc2 = 0.415  ;
KM2 = 43     ;
kc3= 0.02    ;
KM3= 8.3     ;
kc4= 0.02    ;
KM4= 76      ;
kc5= 0.02    ;
KM5= 42      ;
kc6= 0.02    ;
KM6= 10      ;
kc7= 0.01    ;
KM7= 5.1     ;
kc_7=0.89    ;
KM_7=17      ;
kcipp= 4.397 ;
kcdmapp=1    ;
KMipp = 15.6 ;
KMdmapp = 9  ;
kc9= 0.53    ;
KM9=  9.5    ;
kc10= 0.0183 ;
KM10= 7.7    ;
kc11= 0.02   ;
KM11= 125    ;
kc12= 0.002  ;
KM12= 30     ;
kc13= 0.08   ;   %  SMO1
KM13= 500    ;
kc14= 0.05   ;   %  CECI
KM14= 100    ;
kc15= 0.05   ;   %  CYP51
KM15= 160    ;
kc16= 0.05   ;   %  FACKEL
KM16= 100    ;
kc17= 0.05   ;   %  HYDRA
KM17= 100    ;
kc18= 0.005  ;   %  SMT2/3
KM18= 30     ;
kc19= 0.06   ;   %  DWF7 (campesterol branch)
KM19= 140    ;
kc20= 0.06   ;   %  DWF5 (campesterol branch)
KM20= 460    ;
kc21= 0.018  ;   %  DWF1 (campesterol branch)
KM21= 150    ;
% There is neither kc22,nor KM22. Further catalysis of CAMP is not considered.
% (flux into brassinosteroids is very low)
kc23= 0.08   ;   %  SMO2 (sitosterol branch & campesterol branch)
KM23= 480    ;
```

```
kc24= 0.06   ;   %  DWF7 (sitosterol branch)
KM24= 140    ;
kc25= 0.06   ;   %  DWF5 (sitosterol branch)
KM25= 460    ;
kc26= 0.018  ;   %  DWF1 (sitosterol branch)
KM26= 150    ;
kc27= 0.0005 ;   %  CYP710
KM27= 100    ;

kc40= 0.000002 ;   % Conversion of sterol end products into esters and/or
% integration into membranes

kiHMGS = 9  ;       % Competitive inhibition for HMGS by HMGCoA
kiHMGR = 990;       % Competitive inhibition for HMGR by MVA
kiMK   = 0.1;       % Competitive inhibition for MK by FPP
kiSMT1MeCy = 90;    % Competitive inhibition for SMT1 by 24-methylenecycloartanol
kiSMT1Eyloph = 50;  % Competitive inhibition for SMT1 by ethylidenelophenol
kiSMT1sit = 100;    % Competitive inhibition for SMT1 by sitosterol
kiSMT2sit = 300;    % Competitive inhibition for SMT2 by sitosterol

% Enzyme Concentrations [uM]

E1=0.0583;      %AACT
E2=0.0683;      %HMGS
E3=2.3451;      %HMGR
E4=0.0265;      %MK
E5=0.2530;      %PMK
E6=0.0459;      %MPDC
E7=0.1506;      %IPPI
E8=0.1501;      %FPPS
E9=0.0605;      %SQS
E10=0.1396;     %SQE
E11=0.1389;     %CAS
E12=0.17683;    %SMT1 (main pathway)
E13=0.11107;    %SMO1 (main pathway)
E14=0.0209;     %CECI (main pathway)
E15=0.09899;    %CYP51 (main pathway)
E16=0.04549;    %FACKEL (main pathway)
E17=0.1325;     %HYDRA (main pathway)
E18=0.2500;     %SMT2 For Ethyllophenol biosynthesis
E19=0.05452;    %DWF7 (campesterol branch)
E20=0.1615;     %DWF5 (campesterol branch)
E21=0.27268;    %DWF1 (campesterol branch)
%There is no E22. Further catalysis of CAMP is not considered.
E23=0.06996;    %SMO2 (sitosterol branch & campesterol branch)
E24=0.05452;    %DWF7 (sitosterol branch)
E25=0.1615;     %DWF5 (sitosterol branch)
E26=0.27268;    %DWF1 (sitosterol branch)
E27=0.01174;    %CYP710A (sitosterol branch)

% Species equations

xdot=[(kc1*E1*x(1)/(x(1)+KM1)-kc1*E1*x(1)/(x(1)+KM1));% Variation of AcCoA
      kc1*E1*x(1)/(x(1)+KM1)-(kc2*E2*x(2)/KM2)/(1+x(2)/KM2+x(3)/kiHMGS); %Variation of AcAcCoA
      (kc2*E2*x(2)/KM2)/(1+x(2)/KM2+x(3)/kiHMGS)-(E3*kc3*x(3)/KM3)/(1+x(3)/KM3 + x(4)/kiHMGR);
%Variation of HMGCoA
      (E3*kc3*x(3)/KM3)/(1+ x(3)/KM3+ x(4)/kiHMGR)- (E4*kc4*x(4)/KM4)/(1+ x(4)/KM4 + x(9)/kiMK);
%Variation of MVA
      (E4*kc4*x(4)/KM4)/(1+ x(4)/KM4 + x(9)/kiMK)- kc5*E5*x(5)/(x(5)+KM5); %Variation of MVAP
- 5 -
      kc5*E5*x(5)/(x(5)+KM5)-kc6*E6*x(6)/(x(6)+KM6); %Variation of MVAPP
      kc6*E6*x(6)/(x(6)+KM6)+kc_7*E7*x(8)/(x(8)+KM_7)-kc7*E7*x(7)/(x(7)+KM7)-
(kcipp*E8*KMdmapp*x(7)+kcdmapp*E8*KMipp*x(8))/(KMdmapp*x(7)+KMipp*x(8)+KMipp*KMdmapp); %Variation
of IPP
      kc7*E7*x(7)/(x(7)+KM7)-kc_7*E7*x(8)/(x(8)+KM_7)-
(kcipp*E8*KMdmapp*x(7)+kcdmapp*E8*KMipp*x(8))/(KMdmapp*x(7)+KMipp*x(8)+KMipp*KMdmapp); %Variation
of DMAPP
      0.89*(kcipp*E8*KMdmapp*x(7)+kcdmapp*E8*KMipp*x(8))/(KMdmapp*x(7)+KMipp*x(8)+KMipp*KMdmapp)-
kc9*E9*x(9)/(x(9)+KM9); %Variation of FPP - 9 -
      kc9*E9*x(9)/(x(9)+KM9)-kc10*E10*x(10)/(x(10)+KM10); %Variation of Squalene   - 10 -
```

```
      0.97*(kc10*E10*x(10)/(x(10)+KM10))-kc11*E11*x(11)/(x(11)+KM11); %Variation of OxSqualene 
0.97
      kc11*E11*x(11)/(x(11)+KM11)-(kc12*E12*x(12))/(KM12*(1 + x(27)/kiSMT1sit + x(13)/kiSMT1MeCy) 
+x(23)/kiSMT1Eyloph + x(12));  % Variation of Cycloartenol   -12 -
      (kc12*E12*x(12))/(KM12*(1 + x(27)/kiSMT1sit + x(13)/kiSMT1MeCy) +x(23)/kiSMT1Eyloph + 
x(12))- kc13*E13*x(13)/(x(13)+KM13); %Variation in 24-MeCycloartenol - 13 -
      kc13*E13*x(13)/(x(13)+KM13)-kc14*E14*x(14)/(x(14)+KM14); %Variation in Cycloeuclenol - 14 -
      kc14*E14*x(14)/(x(14)+KM14)-kc15*E15*x(15)/(x(15)+KM15); %Variation in Obtusifoliol  - 15 -
      kc15*E15*x(15)/(x(15)+KM15)-kc16*E16*x(16)/(x(16)+KM16); %Variation in CYP51 product  - 16 
-
      kc16*E16*x(16)/(x(16)+KM16)-kc17*E17*x(17)/(x(17)+KM17); %Variation in 4-MeFecosterol  - 17 
-
      kc17*E17*x(17)/(x(17)+KM17)-(kc23*E23*x(18)/(x(18)+KM23))-(kc18*E18*x(18)/KM18)/(1 + 
x(18)/KM18 + x(27)/kiSMT2sit); %Variation in 4-MyLophenol
      (kc23*E23*x(18)/(x(18)+KM23))- (kc19*E19*x(19)/(x(19)+KM19)); %Variation in Episterol
     (kc19*E19*x(19)/(x(19)+KM19))-kc20*E20*x(20)/(x(20)+KM20); %Variation in 5-Dehydroepisterol 
-  26 -
      kc20*E20*x(20)/(x(20)+KM20)-kc21*E21*x(21)/(x(21)+KM21); %Variation in 24-MyCholesterol
      kc21*E21*x(21)/(x(21)+KM21)- kc40*x(22); %Variation in Campesterol  - 28 -
      (kc18*E18*x(18)/KM18)/(1 + x(18)/KM18 + x(27)/kiSMT2sit)-kc23*E23*x(23)/(x(23)+KM23); 
%Variation in 24-EyLophenol - 19 -
      kc23*E23*x(23)/(x(23)+KM23)-(kc24*E24*x(24)/(x(24)+KM24)); %Variation in Avenasterol
     (kc24*E24*x(24)/(x(24)+KM24))-kc25*E25*x(25)/(x(25)+KM25); %Variation in 5-
Dehydroavenasterol   - 21 -
      kc25*E25*x(25)/(x(25)+KM25)-(kc26*E26*x(26)/(x(26)+KM26)); %Variation in Isofucosterol
      (kc26*E26*x(26)/(x(26)+KM26))-kc27*E27*x(27)/(x(27)+KM27) - kc40*x(27); %********Variation 
in Sitosterol - 23 -
      kc27*E27*x(27)/(x(27)+KM27)- kc40*x(28)]; %*******Variation in Stigmasterol  - 24 -
```

**Script file – stem – HMG1**

```
%This program calculates the sterol accumulation patterns in Arabidopsis thaliana

%Definition of Variables
%[AcCoA]=x(1), [AcAcCoA]=x(2) , [HMGCoA]=x(3), [MVA]=x(4), [MVAP]=x(5), [MVAPP]=x(6)
%[IPP]=x(7) , [DMAPP]=x(8), [FPP]=x(9),  [SQ]=x(10),  [OxSQ]=x(11), [CyA]=x(12)
%[MeCyA]=x(13), {...

%clear
xdot = zeros(28,1); %Metabolite vector

%Time interval
tspan = [0 4320000];  %[S]  (50 days)

%vector of initial conditions
xdot0 = [0.065;0;0;0;0;0;0;0;0;0;0;0;0;0;0;0;0;0;0;0;0;0;0;0;0;0;0;0];

%[t,x] = ode15s('Stem_HMG1', tspan, xdot0);
[t,x] = ode15s('Stem_HMG1', tspan, xdot0);

AcCoA=x(:,1);
AcAcCoA=x(:,2);
HMGCoA=x(:,3);
MVA=x(:,4);
MVAP=x(:,5);
MVAPP=x(:,6);
IPP=x(:,7);
DMAPP=x(:,8);
FPP=x(:,9);
SQ=x(:,10);
OxSQ=x(:,11);
CyA=x(:,12);
MeCyA=x(:,13);
Cycloeucalenol=x(:,14);
Obtusifoliol=x(:,15);
UNKNOWN=x(:,16);
MeFecosterol=x(:,17);
MyLophenol=x(:,18);
Episterol=x(:,19);
Dehydroepisterol=x(:,20);
```

```
MyChl=x(:,21);
CAM=x(:,22);
EyLophenol=x(:,23);
Avenasterol=x(:,24);
Dehydroavenasterol=x(:,25);
IFUC=x(:,26);
SIT=x(:,27);
STGM=x(:,28);

%PLOTS

%plot(t/86400,SIT,'b',t/86400,IFUC,'k',t/86400,CyA,'r',t/86400,MeCyA,'c')
plot(t/86400,CyA,'g',t/86400,MeCyA,'k')
hold on
plot(t/86400,CAM,'r','LineWidth',4)
hold on
plot(t/86400,IFUC,'c')
hold on
plot(t/86400,SIT,'b','LineWidth',4)
hold on
plot(t/86400,STGM,'m','LineWidth',4)

%plot(t/86400,SIT,'b',t/86400,IFUC,'k',t/86400,CyA,'r',t/86400,MeCyA,'c',t/86400,OxSQ,'g',
t/86400,CAM,'m', t/86400,AcCoA,'y',t/86400,STGM,'r*')
%plot(t/86400,SIT,'b',t/86400,IFUC,'k',t/86400,CyA,'r',t/86400,MeCyA,'c',t/86400,MyLophenol,'g',t
/86400,EyLophenol,'m',t/86400,MVA,'y',t/86400,OxSQ,'m',t/86400,Cycloeucalenol,'g',t/86400,Obtusif
oliol,'k',t/86400,MeFecosterol,'r',t/86400,Avenasterol,'g',t/86400,Dehydroavenasterol,'g',t/86400
,AcCoA,'g',t/86400,IPP,'r',t/86400,SQ,'r',t/86400,HMGCoA,'g',t/86400,DMAPP,'m')

%legend('SIT','IFUC','CyA','MeCyA','MyLophenol','EyLophenol','MVA','OxSQ') %
legend('CyA','MeCyA','CAM','IFUC','SIT','STGM')

title('Sterol accumulation in Arabidopsis thaliana leaves - HMG1 ')
xlabel('Time (d)')
ylabel('Sterol Concentration (uM/g fresh weight)')
```

**Seed model – wild-type**

```
function xdot = Seed_WT(t,x,E)%it has a flag between x and E

% This function calculaes sterol accumulation in Arabidopsis thaliana
% Organ: Seeds; Genotype: wild-type
% Michaelis-Menten kinetics are assumed to operate, with the following
% exceptions:

%HMGS is Competitively Inhibited by HMGCoA.
%HMGR is Competitively Inhibited by MVA.
%MK is Competitively Inhibited by FPP.
%SMT1 is Competitively Inhibited by SIT and MeCyA.
%SMT2 is Competitively Inhibited by SIT.

% Kinetic Parameters
%(for details see Table 2 of main manuscript)

% kc units: [1/s]
% KM units: [uM]

kc1 = 2.1    ;
KM1 = 770    ;
kc2 = 0.415  ;
KM2 = 43     ;
kc3= 0.02    ;
KM3= 8.3     ;
kc4= 0.02    ;
KM4= 76      ;
kc5= 0.02    ;
KM5= 42      ;
kc6= 0.02    ;
KM6= 10      ;
kc7= 0.01    ;
KM7= 5.1     ;
kc_7=0.89    ;
KM_7=17      ;
kcipp= 4.397 ;
kcdmapp=1    ;
KMipp = 15.6 ;
KMdmapp = 9  ;
kc9= 0.53    ;
KM9=  9.5    ;
kc10= 0.0183 ;
KM10= 7.7    ;
kc11= 0.02   ;
KM11= 125    ;
kc12= 0.002  ;
KM12= 30     ;
kc13= 0.08   ;   %  SMO1
KM13= 500    ;
kc14= 0.05   ;   %  CECI
KM14= 100    ;
kc15= 0.05   ;   %  CYP51
KM15= 160    ;
kc16= 0.05   ;   %  FACKEL
KM16= 100    ;
kc17= 0.05   ;   %  HYDRA
KM17= 100    ;
kc18= 0.005  ;   %  SMT2/3
KM18= 30     ;
kc19= 0.06   ;   %  DWF7 (campesterol branch)
KM19= 140    ;
kc20= 0.06   ;   %  DWF5 (campesterol branch)
KM20= 460    ;
kc21= 0.018  ;   %  DWF1 (campesterol branch)
KM21= 150    ;
% There is neither kc22,nor KM22. Further catalysis of CAMP is not considered.
% (flux into brassinosteroids is very low)
kc23= 0.08   ;   %  SMO2 (sitosterol branch & campesterol branch)
KM23= 480    ;
```

```
kc24= 0.06   ;   %  DWF7 (sitosterol branch)
KM24= 140    ;
kc25= 0.06   ;   %  DWF5 (sitosterol branch)
KM25= 460    ;
kc26= 0.018  ;   %  DWF1 (sitosterol branch)
KM26= 150    ;
kc27= 0.0005 ;   %  CYP710
KM27= 100    ;

kc40= 0.0000005 ;   % Conversion of sterol end products into esters and/or
% integration into membranes

kiHMGS = 9  ;       % Competitive inhibition for HMGS by HMGCoA
kiHMGR = 990;       % Competitive inhibition for HMGR by MVA
kiMK   = 0.1;       % Competitive inhibition for MK by FPP
kiSMT1MeCy = 90;    % Competitive inhibition for SMT1 by 24-methylenecycloartanol
kiSMT1Eyloph = 50;  % Competitive inhibition for SMT1 by ethylidenelophenol
kiSMT1sit = 100;    % Competitive inhibition for SMT1 by sitosterol
kiSMT2sit = 300;    % Competitive inhibition for SMT2 by sitosterol

% Enzyme Concentrations [uM]

E1=0.0583;      %AACT
E2=0.0683;      %HMGS
E3=0.1581;      %HMGR
E4=0.0265;      %MK
E5=0.2530;      %PMK
E6=0.0459;      %MPDC
E7=0.1506;      %IPPI
E8=0.1501;      %FPPS
E9=0.0605;      %SQS
E10=0.1396;     %SQE
E11=0.1389;     %CAS
E12=0.17683;    %SMT1 (main pathway)
E13=0.11107;    %SMO1 (main pathway)
E14=0.0209;     %CECI (main pathway)
E15=0.09899;    %CYP51 (main pathway)
E16=0.04549;    %FACKEL (main pathway)
E17=0.1325;     %HYDRA (main pathway)
E18=0.3251;     %SMT2 For Ethyllophenol biosynthesis
E19=0.05452;    %DWF7 (campesterol branch)
E20=0.1615;     %DWF5 (campesterol branch)
E21=0.27268;    %DWF1 (campesterol branch)
%There is no E22. Further catalysis of CAMP is not considered.
E23=0.06996;    %SMO2 (sitosterol branch & campesterol branch)
E24=0.05452;    %DWF7 (sitosterol branch)
E25=0.1615;     %DWF5 (sitosterol branch)
E26=0.27268;    %DWF1 (sitosterol branch)
E27=0.01174;    %CYP710A (sitosterol branch)

% Species equations

xdot=[(kc1*E1*x(1)/(x(1)+KM1)-kc1*E1*x(1)/(x(1)+KM1));% Variation of AcCoA
      kc1*E1*x(1)/(x(1)+KM1)-(kc2*E2*x(2)/KM2)/(1+x(2)/KM2+x(3)/kiHMGS); %Variation of AcAcCoA
      (kc2*E2*x(2)/KM2)/(1+x(2)/KM2+x(3)/kiHMGS)-(E3*kc3*x(3)/KM3)/(1+x(3)/KM3 + x(4)/kiHMGR);
%Variation of HMGCoA
      (E3*kc3*x(3)/KM3)/(1+ x(3)/KM3+ x(4)/kiHMGR)- (E4*kc4*x(4)/KM4)/(1+ x(4)/KM4 + x(9)/kiMK);
%Variation of MVA
      (E4*kc4*x(4)/KM4)/(1+ x(4)/KM4 + x(9)/kiMK)- kc5*E5*x(5)/(x(5)+KM5); %Variation of MVAP
- 5 -
      kc5*E5*x(5)/(x(5)+KM5)-kc6*E6*x(6)/(x(6)+KM6); %Variation of MVAPP
      kc6*E6*x(6)/(x(6)+KM6)+kc_7*E7*x(8)/(x(8)+KM_7)-kc7*E7*x(7)/(x(7)+KM7)-
(kcipp*E8*KMdmapp*x(7)+kcdmapp*E8*KMipp*x(8))/(KMdmapp*x(7)+KMipp*x(8)+KMipp*KMdmapp); %Variation
of IPP
      kc7*E7*x(7)/(x(7)+KM7)-kc_7*E7*x(8)/(x(8)+KM_7)-
(kcipp*E8*KMdmapp*x(7)+kcdmapp*E8*KMipp*x(8))/(KMdmapp*x(7)+KMipp*x(8)+KMipp*KMdmapp); %Variation
of DMAPP
      0.89*(kcipp*E8*KMdmapp*x(7)+kcdmapp*E8*KMipp*x(8))/(KMdmapp*x(7)+KMipp*x(8)+KMipp*KMdmapp)-
kc9*E9*x(9)/(x(9)+KM9); %Variation of FPP - 9 -
      kc9*E9*x(9)/(x(9)+KM9)-kc10*E10*x(10)/(x(10)+KM10); %Variation of Squalene   - 10 -
```

```
      0.97*(kc10*E10*x(10)/(x(10)+KM10))-kc11*E11*x(11)/(x(11)+KM11); %Variation of OxSqualene 
0.97
      kc11*E11*x(11)/(x(11)+KM11)-(kc12*E12*x(12))/(KM12*(1 + x(27)/kiSMT1sit + x(13)/kiSMT1MeCy) 
+x(23)/kiSMT1Eyloph + x(12));  % Variation of Cycloartenol   -12 -
      (kc12*E12*x(12))/(KM12*(1 + x(27)/kiSMT1sit + x(13)/kiSMT1MeCy) +x(23)/kiSMT1Eyloph + 
x(12))- kc13*E13*x(13)/(x(13)+KM13); %Variation in 24-MeCycloartenol - 13 -
      kc13*E13*x(13)/(x(13)+KM13)-kc14*E14*x(14)/(x(14)+KM14); %Variation in Cycloeuclenol - 14 -
      kc14*E14*x(14)/(x(14)+KM14)-kc15*E15*x(15)/(x(15)+KM15); %Variation in Obtusifoliol  - 15 -
      kc15*E15*x(15)/(x(15)+KM15)-kc16*E16*x(16)/(x(16)+KM16); %Variation in CYP51 product  - 16 
-
      kc16*E16*x(16)/(x(16)+KM16)-kc17*E17*x(17)/(x(17)+KM17); %Variation in 4-MeFecosterol  - 17 
-
      kc17*E17*x(17)/(x(17)+KM17)-(kc23*E23*x(18)/(x(18)+KM23))-(kc18*E18*x(18)/KM18)/(1 + 
x(18)/KM18 + x(27)/kiSMT2sit); %Variation in 4-MyLophenol
      (kc23*E23*x(18)/(x(18)+KM23))- (kc19*E19*x(19)/(x(19)+KM19)); %Variation in Episterol
     (kc19*E19*x(19)/(x(19)+KM19))-kc20*E20*x(20)/(x(20)+KM20); %Variation in 5-Dehydroepisterol 
-  26 -
      kc20*E20*x(20)/(x(20)+KM20)-kc21*E21*x(21)/(x(21)+KM21); %Variation in 24-MyCholesterol
      kc21*E21*x(21)/(x(21)+KM21)- kc40*x(22); %Variation in Campesterol  - 28 -
      (kc18*E18*x(18)/KM18)/(1 + x(18)/KM18 + x(27)/kiSMT2sit)-kc23*E23*x(23)/(x(23)+KM23); 
%Variation in 24-EyLophenol - 19 -
      kc23*E23*x(23)/(x(23)+KM23)-(kc24*E24*x(24)/(x(24)+KM24)); %Variation in Avenasterol
     (kc24*E24*x(24)/(x(24)+KM24))-kc25*E25*x(25)/(x(25)+KM25); %Variation in 5-
Dehydroavenasterol   - 21 -
      kc25*E25*x(25)/(x(25)+KM25)-(kc26*E26*x(26)/(x(26)+KM26)); %Variation in Isofucosterol
      (kc26*E26*x(26)/(x(26)+KM26))-kc27*E27*x(27)/(x(27)+KM27) - kc40*x(27); %********Variation 
in Sitosterol - 23 -
      kc27*E27*x(27)/(x(27)+KM27)- kc40*x(28)]; %*******Variation in Stigmasterol  - 24 -
```

**Script file – seed – wild-type**

```
%This program calculates the sterol accumulation patterns in Arabidopsis thaliana

%Definition of Variables
%[AcCoA]=x(1), [AcAcCoA]=x(2) , [HMGCoA]=x(3), [MVA]=x(4), [MVAP]=x(5), [MVAPP]=x(6)
%[IPP]=x(7) , [DMAPP]=x(8), [FPP]=x(9),  [SQ]=x(10),  [OxSQ]=x(11), [CyA]=x(12)
%[MeCyA]=x(13), {...

%clear
xdot = zeros(28,1); %Metabolite vector

%Time interval
tspan = [0 4320000];  %[S]  (50 days)

%vector of initial conditions
xdot0 = [0.065;0;0;0;0;0;0;0;0;0;0;0;0;0;0;0;0;0;0;0;0;0;0;0;0;0;0;0];

%[t,x] = ode15s('Seed_WT', tspan, xdot0);
[t,x] = ode15s('Seed_WT', tspan, xdot0);

AcCoA=x(:,1);
AcAcCoA=x(:,2);
HMGCoA=x(:,3);
MVA=x(:,4);
MVAP=x(:,5);
MVAPP=x(:,6);
IPP=x(:,7);
DMAPP=x(:,8);
FPP=x(:,9);
SQ=x(:,10);
OxSQ=x(:,11);
CyA=x(:,12);
MeCyA=x(:,13);
Cycloeucalenol=x(:,14);
Obtusifoliol=x(:,15);
UNKNOWN=x(:,16);
MeFecosterol=x(:,17);
MyLophenol=x(:,18);
Episterol=x(:,19);
Dehydroepisterol=x(:,20);
```

```
MyChl=x(:,21);
CAM=x(:,22);
EyLophenol=x(:,23);
Avenasterol=x(:,24);
Dehydroavenasterol=x(:,25);
IFUC=x(:,26);
SIT=x(:,27);
STGM=x(:,28);

%PLOTS

%plot(t/86400,SIT,'b',t/86400,IFUC,'k',t/86400,CyA,'r',t/86400,MeCyA,'c')
plot(t/86400,CyA,'g',t/86400,MeCyA,'k')
hold on
plot(t/86400,CAM,'r','LineWidth',4)
hold on
plot(t/86400,IFUC,'c')
hold on
plot(t/86400,SIT,'b','LineWidth',4)
hold on
plot(t/86400,STGM,'m','LineWidth',4)

%plot(t/86400,SIT,'b',t/86400,IFUC,'k',t/86400,CyA,'r',t/86400,MeCyA,'c',t/86400,OxSQ,'g',
t/86400,CAM,'m', t/86400,AcCoA,'y',t/86400,STGM,'r*')
%plot(t/86400,SIT,'b',t/86400,IFUC,'k',t/86400,CyA,'r',t/86400,MeCyA,'c',t/86400,MyLophenol,'g',t
/86400,EyLophenol,'m',t/86400,MVA,'y',t/86400,OxSQ,'m',t/86400,Cycloeucalenol,'g',t/86400,Obtusif
oliol,'k',t/86400,MeFecosterol,'r',t/86400,Avenasterol,'g',t/86400,Dehydroavenasterol,'g',t/86400
,AcCoA,'g',t/86400,IPP,'r',t/86400,SQ,'r',t/86400,HMGCoA,'g',t/86400,DMAPP,'m')

%legend('SIT','IFUC','CyA','MeCyA','MyLophenol','EyLophenol','MVA','OxSQ') %
legend('CyA','MeCyA','CAM','IFUC','SIT','STGM')

title('Sterol accumulation in Arabidopsis thaliana leaves - HMG1 ')
xlabel('Time (d)')
ylabel('Sterol Concentration (uM/g fresh weight)')
```

**Seed model – HMG1**

```
function xdot = Seed_HMG1(t,x,E)%it has a flag between x and E

% This function calculaes sterol accumulation in Arabidopsis thaliana
% Organ: Seeds; Genotype: HMG1
% Michaelis-Menten kinetics are assumed to operate, with the following
% exceptions:

%HMGS is Competitively Inhibited by HMGCoA.
%HMGR is Competitively Inhibited by MVA.
%MK is Competitively Inhibited by FPP.
%SMT1 is Competitively Inhibited by SIT and MeCyA.
%SMT2 is Competitively Inhibited by SIT.

% Kinetic Parameters
%(for details see Table 2 of main manuscript)

% kc units: [1/s]
% KM units: [uM]

kc1 = 2.1    ;
KM1 = 770    ;
kc2 = 0.415  ;
KM2 = 43     ;
kc3= 0.02    ;
KM3= 8.3     ;
kc4= 0.02    ;
KM4= 76      ;
kc5= 0.02    ;
KM5= 42      ;
kc6= 0.02    ;
KM6= 10      ;
kc7= 0.01    ;
KM7= 5.1     ;
kc_7=0.89    ;
KM_7=17      ;
kcipp= 4.397 ;
kcdmapp=1    ;
KMipp = 15.6 ;
KMdmapp = 9  ;
kc9= 0.53    ;
KM9=  9.5    ;
kc10= 0.0183 ;
KM10= 7.7    ;
kc11= 0.02   ;
KM11= 125    ;
kc12= 0.002  ;
KM12= 30     ;
kc13= 0.08   ;   %  SMO1
KM13= 500    ;
kc14= 0.05   ;   %  CECI
KM14= 100    ;
kc15= 0.05   ;   %  CYP51
KM15= 160    ;
kc16= 0.05   ;   %  FACKEL
KM16= 100    ;
kc17= 0.05   ;   %  HYDRA
KM17= 100    ;
kc18= 0.005  ;   %  SMT2/3
KM18= 30     ;
kc19= 0.06   ;   %  DWF7 (campesterol branch)
KM19= 140    ;
kc20= 0.06   ;   %  DWF5 (campesterol branch)
KM20= 460    ;
kc21= 0.018  ;   %  DWF1 (campesterol branch)
KM21= 150    ;
% There is neither kc22,nor KM22. Further catalysis of CAMP is not considered.
% (flux into brassinosteroids is very low)
kc23= 0.08   ;   %  SMO2 (sitosterol branch & campesterol branch)
KM23= 480    ;
```

```
kc24= 0.06   ;   %  DWF7 (sitosterol branch)
KM24= 140    ;
kc25= 0.06   ;   %  DWF5 (sitosterol branch)
KM25= 460    ;
kc26= 0.018  ;   %  DWF1 (sitosterol branch)
KM26= 150    ;
kc27= 0.0005 ;   %  CYP710
KM27= 100    ;

kc40= 0.000002 ;   % Conversion of sterol end products into esters and/or
% integration into membranes

kiHMGS = 9  ;        % Competitive inhibition for HMGS by HMGCoA
kiHMGR = 990;        % Competitive inhibition for HMGR by MVA
kiMK   = 0.1;        % Competitive inhibition for MK by FPP
kiSMT1MeCy = 90;     % Competitive inhibition for SMT1 by 24-methylenecycloartanol
kiSMT1Eyloph = 50;   % Competitive inhibition for SMT1 by ethylidenelophenol
kiSMT1sit = 100;     % Competitive inhibition for SMT1 by sitosterol
kiSMT2sit = 300;     % Competitive inhibition for SMT2 by sitosterol

% Enzyme Concentrations [uM]

E1=0.0583;      %AACT
E2=0.0683;      %HMGS
E3=0.3162;      %HMGR
E4=0.0265;      %MK
E5=0.2530;      %PMK
E6=0.0459;      %MPDC
E7=0.1506;      %IPPI
E8=0.1501;      %FPPS
E9=0.0605;      %SQS
E10=0.1396;     %SQE
E11=0.1389;     %CAS
E12=0.17683;    %SMT1 (main pathway)
E13=0.11107;    %SMO1 (main pathway)
E14=0.0209;     %CECI (main pathway)
E15=0.09899;    %CYP51 (main pathway)
E16=0.04549;    %FACKEL (main pathway)
E17=0.1325;     %HYDRA (main pathway)
E18=0.3251;     %SMT2 For Ethyllophenol biosynthesis
E19=0.05452;    %DWF7 (campesterol branch)
E20=0.1615;     %DWF5 (campesterol branch)
E21=0.27268;    %DWF1 (campesterol branch)
%There is no E22. Further catalysis of CAMP is not considered.
E23=0.06996;    %SMO2 (sitosterol branch & campesterol branch)
E24=0.05452;    %DWF7 (sitosterol branch)
E25=0.1615;     %DWF5 (sitosterol branch)
E26=0.27268;    %DWF1 (sitosterol branch)
E27=0.01174;    %CYP710A (sitosterol branch)

% Species equations

xdot=[(kc1*E1*x(1)/(x(1)+KM1)-kc1*E1*x(1)/(x(1)+KM1));% Variation of AcCoA
      kc1*E1*x(1)/(x(1)+KM1)-(kc2*E2*x(2)/KM2)/(1+x(2)/KM2+x(3)/kiHMGS); %Variation of AcAcCoA
      (kc2*E2*x(2)/KM2)/(1+x(2)/KM2+x(3)/kiHMGS)-(E3*kc3*x(3)/KM3)/(1+x(3)/KM3 + x(4)/kiHMGR);
%Variation of HMGCoA
      (E3*kc3*x(3)/KM3)/(1+ x(3)/KM3+ x(4)/kiHMGR)- (E4*kc4*x(4)/KM4)/(1+ x(4)/KM4 + x(9)/kiMK);
%Variation of MVA
      (E4*kc4*x(4)/KM4)/(1+ x(4)/KM4 + x(9)/kiMK)- kc5*E5*x(5)/(x(5)+KM5); %Variation of MVAP
- 5 -
      kc5*E5*x(5)/(x(5)+KM5)-kc6*E6*x(6)/(x(6)+KM6); %Variation of MVAPP
      kc6*E6*x(6)/(x(6)+KM6)+kc_7*E7*x(8)/(x(8)+KM_7)-kc7*E7*x(7)/(x(7)+KM7)-
(kcipp*E8*KMdmapp*x(7)+kcdmapp*E8*KMipp*x(8))/(KMdmapp*x(7)+KMipp*x(8)+KMipp*KMdmapp); %Variation
of IPP
      kc7*E7*x(7)/(x(7)+KM7)-kc_7*E7*x(8)/(x(8)+KM_7)-
(kcipp*E8*KMdmapp*x(7)+kcdmapp*E8*KMipp*x(8))/(KMdmapp*x(7)+KMipp*x(8)+KMipp*KMdmapp); %Variation
of DMAPP
      0.89*(kcipp*E8*KMdmapp*x(7)+kcdmapp*E8*KMipp*x(8))/(KMdmapp*x(7)+KMipp*x(8)+KMipp*KMdmapp)-
kc9*E9*x(9)/(x(9)+KM9); %Variation of FPP - 9 -
      kc9*E9*x(9)/(x(9)+KM9)-kc10*E10*x(10)/(x(10)+KM10); %Variation of Squalene   - 10 -
```

```
      0.97*(kc10*E10*x(10)/(x(10)+KM10))-kc11*E11*x(11)/(x(11)+KM11); %Variation of OxSqualene
0.97
      kc11*E11*x(11)/(x(11)+KM11)-(kc12*E12*x(12))/(KM12*(1 + x(27)/kiSMT1sit + x(13)/kiSMT1MeCy)
+x(23)/kiSMT1Eyloph + x(12));  % Variation of Cycloartenol   -12 -
      (kc12*E12*x(12))/(KM12*(1 + x(27)/kiSMT1sit + x(13)/kiSMT1MeCy) +x(23)/kiSMT1Eyloph +
x(12))- kc13*E13*x(13)/(x(13)+KM13); %Variation in 24-MeCycloartenol - 13 -
      kc13*E13*x(13)/(x(13)+KM13)-kc14*E14*x(14)/(x(14)+KM14); %Variation in Cycloeuclenol - 14 -
      kc14*E14*x(14)/(x(14)+KM14)-kc15*E15*x(15)/(x(15)+KM15); %Variation in Obtusifoliol  - 15 -
      kc15*E15*x(15)/(x(15)+KM15)-kc16*E16*x(16)/(x(16)+KM16); %Variation in CYP51 product  - 16
-
      kc16*E16*x(16)/(x(16)+KM16)-kc17*E17*x(17)/(x(17)+KM17); %Variation in 4-MeFecosterol  - 17
-
      kc17*E17*x(17)/(x(17)+KM17)-(kc23*E23*x(18)/(x(18)+KM23))-(kc18*E18*x(18)/KM18)/(1 +
x(18)/KM18 + x(27)/kiSMT2sit); %Variation in 4-MyLophenol
      (kc23*E23*x(18)/(x(18)+KM23))- (kc19*E19*x(19)/(x(19)+KM19)); %Variation in Episterol
     (kc19*E19*x(19)/(x(19)+KM19))-kc20*E20*x(20)/(x(20)+KM20); %Variation in 5-Dehydroepisterol
-  26 -
      kc20*E20*x(20)/(x(20)+KM20)-kc21*E21*x(21)/(x(21)+KM21); %Variation in 24-MyCholesterol
      kc21*E21*x(21)/(x(21)+KM21)- kc40*x(22); %Variation in Campesterol  - 28 -
      (kc18*E18*x(18)/KM18)/(1 + x(18)/KM18 + x(27)/kiSMT2sit)-kc23*E23*x(23)/(x(23)+KM23);
%Variation in 24-EyLophenol - 19 -
      kc23*E23*x(23)/(x(23)+KM23)-(kc24*E24*x(24)/(x(24)+KM24)); %Variation in Avenasterol
     (kc24*E24*x(24)/(x(24)+KM24))-kc25*E25*x(25)/(x(25)+KM25); %Variation in 5-
Dehydroavenasterol   - 21 -
      kc25*E25*x(25)/(x(25)+KM25)-(kc26*E26*x(26)/(x(26)+KM26)); %Variation in Isofucosterol
      (kc26*E26*x(26)/(x(26)+KM26))-kc27*E27*x(27)/(x(27)+KM27) - kc40*x(27); %********Variation
in Sitosterol - 23 -
      kc27*E27*x(27)/(x(27)+KM27)- kc40*x(28)]; %*******Variation in Stigmasterol  - 24 -
```

**Script file – seed- HMG1**

```
%This program calculates the sterol accumulation patterns in Arabidopsis thaliana

%Definition of Variables
%[AcCoA]=x(1), [AcAcCoA]=x(2) , [HMGCoA]=x(3), [MVA]=x(4), [MVAP]=x(5), [MVAPP]=x(6)
%[IPP]=x(7) , [DMAPP]=x(8), [FPP]=x(9),  [SQ]=x(10),  [OxSQ]=x(11), [CyA]=x(12)
%[MeCyA]=x(13), {...

%clear
xdot = zeros(28,1); %Metabolite vector

%Time interval
tspan = [0 4320000];  %[S]  (50 days)

%vector of initial conditions
xdot0 = [0.3;0;0;0;0;0;0;0;0;0;0;0;0;0;0;0;0;0;0;0;0;0;0;0;0;0;0;0];

%[t,x] = ode15s('Seed_HMG1', tspan, xdot0);
[t,x] = ode15s('Seed_HMG1', tspan, xdot0);

AcCoA=x(:,1);
AcAcCoA=x(:,2);
HMGCoA=x(:,3);
MVA=x(:,4);
MVAP=x(:,5);
MVAPP=x(:,6);
IPP=x(:,7);
DMAPP=x(:,8);
FPP=x(:,9);
SQ=x(:,10);
OxSQ=x(:,11);
CyA=x(:,12);
MeCyA=x(:,13);
Cycloeucalenol=x(:,14);
Obtusifoliol=x(:,15);
UNKNOWN=x(:,16);
MeFecosterol=x(:,17);
MyLophenol=x(:,18);
Episterol=x(:,19);
Dehydroepisterol=x(:,20);
```

```
MyChl=x(:,21);
CAM=x(:,22);
EyLophenol=x(:,23);
Avenasterol=x(:,24);
Dehydroavenasterol=x(:,25);
IFUC=x(:,26);
SIT=x(:,27);
STGM=x(:,28);

%PLOTS

%plot(t/86400,SIT,'b',t/86400,IFUC,'k',t/86400,CyA,'r',t/86400,MeCyA,'c')
plot(t/86400,CyA,'g',t/86400,MeCyA,'k')
hold on
plot(t/86400,CAM,'r','LineWidth',4)
hold on
plot(t/86400,IFUC,'c')
hold on
plot(t/86400,SIT,'b','LineWidth',4)
hold on
plot(t/86400,STGM,'m','LineWidth',4)

%plot(t/86400,SIT,'b',t/86400,IFUC,'k',t/86400,CyA,'r',t/86400,MeCyA,'c',t/86400,OxSQ,'g',
t/86400,CAM,'m', t/86400,AcCoA,'y',t/86400,STGM,'r*')
%plot(t/86400,SIT,'b',t/86400,IFUC,'k',t/86400,CyA,'r',t/86400,MeCyA,'c',t/86400,MyLophenol,'g',t
/86400,EyLophenol,'m',t/86400,MVA,'y',t/86400,OxSQ,'m',t/86400,Cycloeucalenol,'g',t/86400,Obtusif
oliol,'k',t/86400,MeFecosterol,'r',t/86400,Avenasterol,'g',t/86400,Dehydroavenasterol,'g',t/86400
,AcCoA,'g',t/86400,IPP,'r',t/86400,SQ,'r',t/86400,HMGCoA,'g',t/86400,DMAPP,'m')

%legend('SIT','IFUC','CyA','MeCyA','MyLophenol','EyLophenol','MVA','OxSQ') %
legend('CyA','MeCyA','CAM','IFUC','SIT','STGM')

title('Sterol accumulation in Arabidopsis thaliana leaves - HMG1 ')
xlabel('Time (d)')
ylabel('Sterol Concentration (uM/g fresh weight)')
```
